# LGTM: Gaussian Process Modulated Neural Topic Modeling for Longitudinal Microbiome

**DOI:** 10.64898/2026.04.10.717451

**Authors:** Xiao Yuan, Ádám Arany, András Formanek, Yves Moreau, Harri Lähdesmäki, Tommi Vatanen

## Abstract

Longitudinal microbiome data are key to understanding the dynamics of microbial communities and their relationships with the host and environment. However, analysis of such data is challenging due to high dimensionality, compositionality, irregular sampling and temporal dependencies on external covariates. Existing analytical approaches typically address only subsets of these challenges, limiting their ability to yield biologically interpretable insights. We introduce LGTM, a probabilistic modeling framework that combines flexible non-linear longitudinal modeling with interpretable topic-based representations of the microbiome. LGTM simultaneously identifies microbial co-abundance patterns (“topics”) and models how their proportions change over time and in relation to host and environmental covariates. Using multiple longitudinal human gut microbiome datasets, we demonstrate that LGTM identifies diverse microbial topics whose major patterns are reproducible across runs, while achieving competitive performance in imputation and forecasting tasks. A key strength of the framework is its interpretability: LGTM yields microbial topics with biologically interpretable taxonomic compositions and directly quantifies associations between covariates and microbial dynamics. LGTM is available at https://github.com/yuanx749/lgtm.

## 1. Introduction

Longitudinal gut microbiome studies provide a window into microbial ecology *in vivo*, enabling direct observation of community assembly, succession, and stability across host development or perturbations [1, 2, 3]. By repeatedly sampling the same individuals over time, these data offer a unique opportunity to characterize the dynamics and individualized ecological trajectories, and their associations with host and environmental covariates [4, 5, 6, 7]. However, analyzing longitudinal microbiome data presents several challenges. First, they are high-dimensional, often comprising hundreds to thousands of microbial taxa. Second, relative abundances are compositional, meaning that they represent proportions constrained to sum to one. Third, longitudinal sampling is often sparse and irregular due to practical constraints common in gut microbiome studies, leading to missing data at various timepoints for many subjects. Fourth, microbial compositions exhibit complex temporal patterns that can vary across subjects and groups, influenced by factors like diet, disease state, medications or other host-associated perturbations. Together, these characteristics substantially complicate inference of meaningful ecological trajectories from longitudinal gut microbiome data using existing analytical approaches.

Existing approaches for analyzing longitudinal microbiome data include linear mixed-effects models (LMMs), generalized Lotka–Volterra (gLV) models, Gaussian processes (GPs) and neural network-based models. Beyond accurate modeling of temporal dynamics, longitudinal gut microbiome studies require interpretable representations that enable comparison of microbial trajectories across individuals and conditions. LMMs are widely used for association analysis [8] but analyze taxa independently and struggle to flexibly capture non-linear temporal patterns or inter-taxa dependencies. The gLV equations model interactions between taxa [9, 10] but they are mechanistic and typically only applicable to simple model communities with a limited number of taxa. GPs offer a non-parametric approach for time-series modeling and have been applied to longitudinal gut microbiome data [11, 12], yet they do not capture interactions between taxa. Neural networks are particularly powerful at modeling high-dimensional data. Deep learning-based sequence models, including recurrent neural networks (RNNs) and transformers, have shown strong performance in imputing general time-series data [13, 14, 15]. While RNN-based models have been applied to microbiome data [16], they require discretizing time into intervals and do not quantify uncertainty. Neural differential equation-based methods [17] have also been explored for microbiome analysis [18], but external covariates beyond time were ignored and taxa selections were performed manually. Generative models, especially variational autoencoders (VAEs), are distinguished by their ability to learn low-dimensional representations and generate imputations with inherent uncertainty [19, 20]. Conditional VAEs with GP priors are introduced to capture shared and individual temporal patterns while also handling auxiliary covariates [21, 22, 23]. Recently, an approach leveraging basis function approximation for GP priors has been proposed to overcome the scalability issue [24], demonstrating state-of-the-art performance on imputation and forecasting tasks on general longitudinal data. Nevertheless, applying these models to gut microbiome data requires explicit treatment of compositionality; moreover, despite their expressiveness, most neural network-based approaches lack direct biological interpretability.

A complementary line of research for high-dimensional data analysis focuses on unsupervised dimensionality reduction. Topic modeling, originating from natural language processing, discovers latent topics from a collection of documents and infers the topics a document contains, where a topic is described as a group of related words [25]. Foundational methods include nonnegative matrix factorization (NMF) [26] and probabilistic graphical models built on latent Dirichlet allocation (LDA) [25]. Classical LDA requires labor-intensive, model-specific derivations for parameter updates. In the deep learning paradigm, neural topic models are proposed for efficient and flexible inference [27, 28, 29]. Because of the interpretability and suitability for count data, topic models have been applied to biological sequencing data. While modern neural topic models are increasingly used in single-cell omics to learn complex gene signatures [30, 31], microbiome researchers have mainly relied on classical methods to reveal microbial subcommunities [32, 33]. This line of work has discussed and advanced latent decomposition methods in microbiome data analysis [34], spanning latent variable modeling [35], longitudinal and perturbation analyses [36], and topic alignment across resolutions [37]. Most topic-modeling approaches, however, are static. They model data at the sample level and do not account for temporal correlations between samples or the influence of covariates. This motivates a neural framework in which differentiable components can be composed with temporal and covariate modules under a unified end-to-end formulation.

To address these challenges, we present Longitudinal Gaussian process modulated neural Topic modeling for Microbiome data analysis (LGTM). By jointly learning latent factors and external covariates, our model yields interpretable representations of gut microbial dynamics, with latent factors characterized by key taxa and linked to host and environmental covariates. Using multiple longitudinal gut microbiome datasets, we demonstrate that LGTM achieves competitive predictive performance while identifying diverse microbial topics, corresponding to latent factors, that co-vary with external covariates in biologically meaningful patterns. Together, these results establish a framework for analyzing longitudinal gut microbiome data that supports biologically interpretable investigation of microbial ecological trajectories over time.

## 2. Methods

### 2.1. Problem Formulation

We consider a longitudinal microbiome study comprising *N* samples collected from multiple subjects across time, each sample profiled for *D* microbial taxa (e.g., species). Let 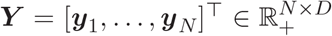 denote the relative abundance table. Here, each row *y* ∈ Δ^*D*^ represents the microbial profile of a sample, and Δ^*D*^ denotes a standard (*D* − 1)-simplex, i.e., the space of nonnegative vectors summing to one. Additionally, each sample is associated with a set of metadata or covariates, collectively denoted as *X* = [*x*_1_, …, *x*_*N*_]^⊤^, where 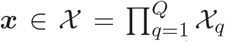, and *X*_*q*_ is the domain of the *q*-th covariate. The covariates may be continuous (e.g., time) or categorical (e.g., subject ID).

Let *L* be the number of topics or latent dimensionality, where *L* ≪ *D*. Each topic is defined as a distribution over the *D* taxa, i.e., topic-taxon distribution *β* ∈ Δ^*D*^. The full set of topics forms the topic matrix 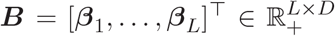. Each sample’s microbial profile is modeled as a mixture of these topics. The topic proportions for all samples are denoted as 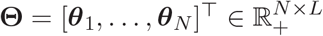, where *θ* ∈ Δ^*L*^ is the sample-topic distribution and also the low-dimensional latent representation of a sample.

Intuitively, LGTM has three coupled parts. The encoder learns sample-level topic mixtures, the decoder learns which taxa characterize the shared topics, and the covariate module learns how the topic mixtures change with time and other metadata. The topics are shared across samples, and covariates affect how the shared topics are mixed in each sample. In the covariate module, each covariate or covariate-time interaction contributes a separate component to the topic proportions. This additive structure allows us to summarize which covariates are associated with variation in each topic. See Figure 1 for an overview of the model.

**Figure 1:**
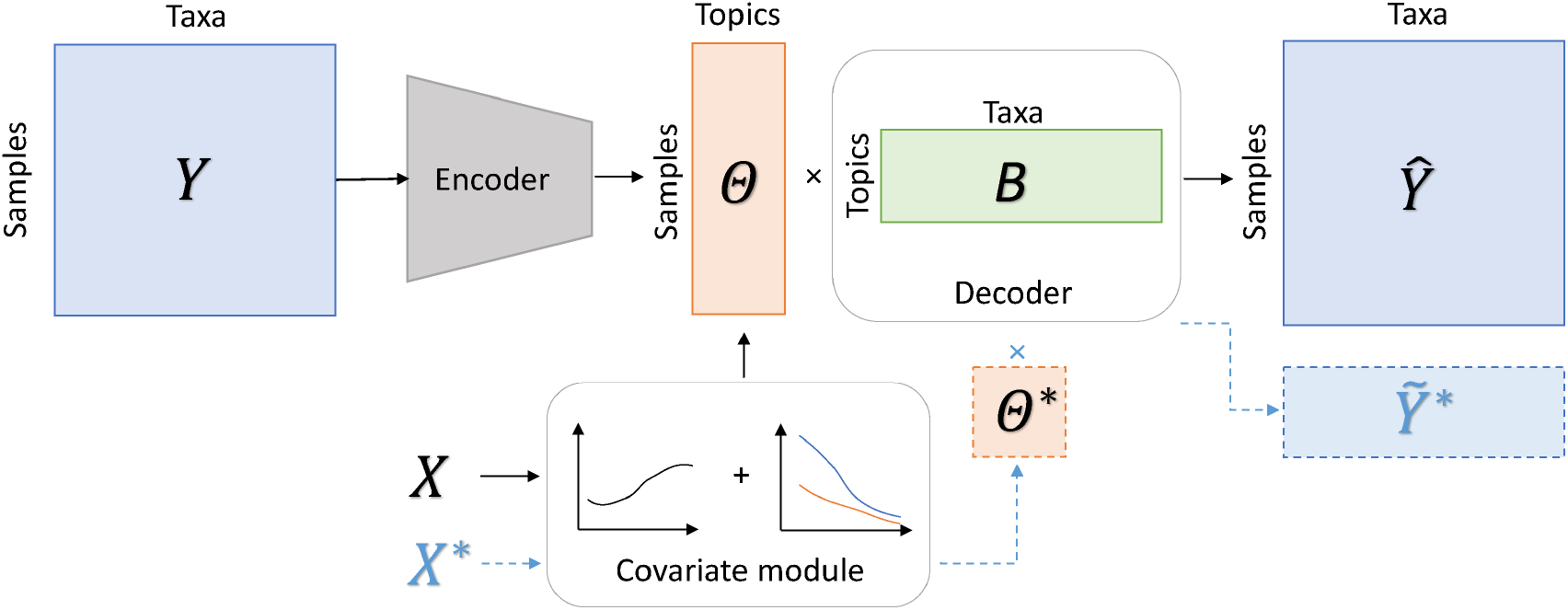
Overview of LGTM. The observed microbiome profile *Y* is mapped by an encoder to topic proportions Θ in a simplex-valued latent space, and decoded by the topic-taxon matrix *B* to reconstruct *Ŷ* . In parallel, a covariate module uses additive GP components via basis-function approximations to map covariates *X* (e.g., time, subject ID, and other metadata) to generate covariate-modulated topic proportions. During training, the encoder and GP-modulated topic proportions are aligned via a KL term, together with cross-entropy reconstruction and GP regularization, and all components are optimized end-to-end. LGTM can perform both topic discovery (*Y* ≈ Θ*B*) and prediction for new covariates using the learned decoder. The dashed pathway illustrates prediction for new covariates *X*^∗^, where predicted topic proportions Θ^∗^ are combined with the shared topic-taxon matrix *B* to generate 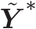.

### 2.2. Gaussian Process and Kernel Approximation

A GP regression model [38] defines a distribution over functions of the form *f* : *X* → ℝ, where is *X* an arbitrary domain. It assumes that function values at any finite set of inputs are jointly Gaussian. Such a process, denoted by *f* (*x*) ∼ *GP* (*µ*(*x*), *K*(*x, x*^′^)), is defined by a mean function *µ*(*x*) (typically zero) and a covariance function *K* (*x, x*^′^), which is any positive definite kernel. The kernel encodes prior knowledge about the function’s properties, such as smoothness or non-linearity.

A primary limitation of GPs is the time complexity that scales as *O* (*N* ^3^) due to the need to invert the kernel matrix. To overcome this, we adopt basis function approaches [39, 40, 24] to approximate the kernels and the GPs. Stationary continuous kernels are covariance functions that depend only on the distance between two inputs, i.e., *K*(*x, x*^′^) = *K*(|*x* − *x*^′^|) = K(*r*), where *x* is a univariate covariate. By Bochner’s theorem, any stationary continuous kernel can be represented as the Fourier transform of a positive measure: 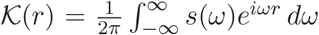, where *s*(*ω*) is the associated spectral density. It has been shown that such stationary kernels can be approximated as [39]:

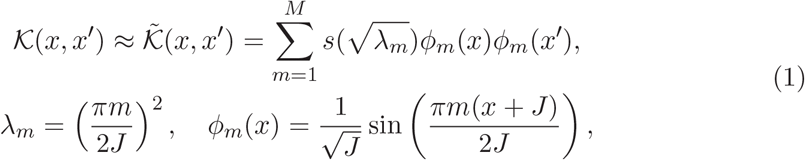

where *λ*_*m*_ and *φ*_*m*_(*x*) are the *m*-th eigenvalue and eigenfunction of the Laplace operator on a bounded interval [− *J, J*]. The Hilbert space approximation allows the GP to be represented in a linear parametric form using basis functions *φ*(*x*) = [*φ*_1_(*x*), …, *φ*_*M*_ (*x*)]^⊤^ ∈ ℝ^*M*^ and corresponding weights *a* ∈ ℝ^*M*^ :

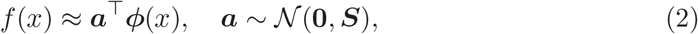

where 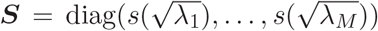. This achieves a time complexity of *O*(*NM*) linear in *N* .

We use the squared exponential (SE) kernel *K*_se_(*x, x*^′^ | *σ, ℓ*) for continuous covariates, where *σ* and *ℓ* denote the amplitude and the length scale, respectively. We also model categorical covariates and their interactions with continuous covariates using basis functions [40]. See Appendix A for details.

### 2.3. Generative Topic Models

We now describe the probabilistic topic model that forms the core of LGTM. Drawing an analogy to natural language documents, each sample is treated as a “document” composed of sequencing reads, and the taxon to which a read maps corresponds to a “word” from a vocabulary of taxa. We assume that each sample can be represented as a mixture of microbial subcommunities, which are referred to as topics. We use “subcommunity” as an operational term for a co-abundance topic, i.e., a statistical association among taxa, not an ecological guild or functional group. Consider a collection of *N* samples with *D* taxa and *L* topics. Topic modeling can be viewed as a non-negative Bayesian matrix factorization. The relative abundance matrix 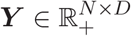 is factorized into two low-rank matrices, 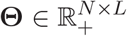 and 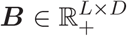, where each row of them lies on a simplex, *θ*_*n*_ ∈ Δ^*L*^ and *β*_*l*_ ∈ Δ^*D*^. We directly model relative abundance, which is the primary quantity of interest in many microbiome analyses, as raw read counts do not reflect the actual microbial load and need to be normalized to account for variations in sequencing depth [41, 42].

Unlike standard LDAs, where the latent variable *θ* follows a Dirichlet [25] or a logistic-normal distribution [27], we condition *θ* on covariates *x*. Specifically, we use GPs to modulate the latent space. GPs can be extended to multiple outputs and multiple additive components [43, 23], in the form *f* ^(*r*)^ :*X*^(*r*)^ →ℝ^*L*^, where *r* ∈ {1, …, *R*} and *R* is the number of components. Each additive GP component depends on a single or a pair of covariates *x*^(*r*)^ ∈ *X*^(*r*)^ ⊆ *X*. The sum of these components is the cumulative effect of the covariates. The additive GP structure allows modeling shared and random effects and is interpretable similarly to LMMs.

Concretely, for our longitudinal data, we construct additive components as follows. One component uses an SE kernel on time to model shared temporal patterns, and one component uses a categorical kernel on subject ID to model individual effects. For each of the other categorical covariates, we add an interaction component using an interaction kernel that depends on time and a categorical covariate, to model group-specific temporal patterns.

Following [24], we apply the Hilbert space approximation to each component and each latent dimension separately. Take the SE kernel as an example, and let 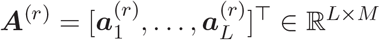 denote the linear parameters from Eq. (2). Following Eqs. (1)–(2), the *r*-th component is approximated as:

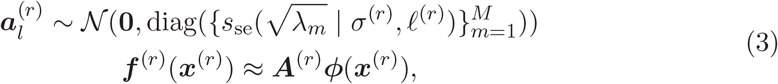

where *s*_se_(·) is the spectral density of the SE kernel, and *σ*^(*r*)^ and *ℓ*^(*r*)^ have log-normal priors. This formulation extends directly to interaction kernels. For enhanced interpretability of the component latent functions, we depart from previous work [23, 24] that sums the components. Instead, we apply a softmax transformation to constrain each component to the simplex, and then average them to obtain the topic proportions:

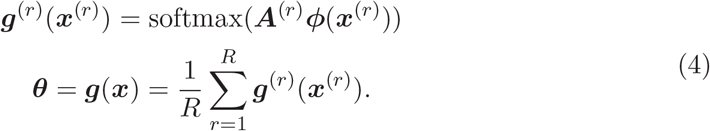

For samples with missing covariates, we ignore components *g*^(*r*)^(*x*^(*r*)^) that have missing covariates and simply average over the components with observed covariates. Consequently, the overall covariate effect on topic proportions is decomposed into functions of individual covariates and their interactions. Finally, the decoder produces the relative abundance from the topic-taxon distribution matrix and the topic proportions: 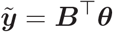.

### 2.4. Training and Optimization

We adopt an autoencoding architecture as in past work on neural topic models. The encoder network maps each sample to latent topic proportions through a logistic-normal transformation (a Gaussian followed by a softmax). The encoder is parameterized by a multilayer perceptron (MLP) with one hidden layer. The decoder is a single linear layer. Each row of its weight matrix is a softmax-transformed vector of learnable parameters. To mitigate instability from random initialization, we apply nonnegative double singular value decomposition (NNDSVD) [44] on the training data and initialize decoder’s parameters based on its output.

During training, the model reconstructs the relative abundance via 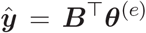, where *θ*^(*e*)^ is sampled from the encoder distribution *q*(*θ*|*y*). In parallel, the covariate module generates GP-modulated topic proportions *θ*^(*g*)^, which are used for prediction via 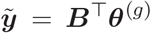. Note that the topic proportions *θ*^(*e*)^ and *θ*^(*g*)^ can be interpreted as probability distributions over topics. The covariate module is parameterized by a collection of weights 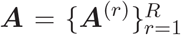 that provide an approximation for the additive GP model in Eq. (4). These weights are endowed with the prior given in Eq. (3). All model parameters are jointly optimized by minimizing a loss composed of a reconstruction term and Kullback–Leibler (KL) regularization terms. The loss for one sample is defined as:

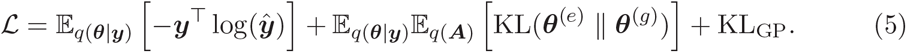

The first term is the cross-entropy between the observation *y* and its reconstruction *ŷ*. Note that under a multinomial assumption, the log-likelihood reduces to cross-entropy up to a constant (Appendix F.2.1). The second term measures the KL divergence between the encoder output and the output from the covariate module across the *L* topics. It is computed as 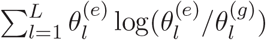, where expectations are approximated using Monte Carlo, and 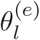 and 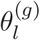 are the *l*-th elements of *θ*^(*e*)^ and *θ*^(*g*)^, respectively. This term closes the gap between the training and testing pipelines, similar to the idea of conditional VAEs [20]. Because the covariate module outputs the mean of softmax-transformed additive GPs and does not yield a closed-form density, this KL term provides a practical way to encourage consistency between the learned latent representation (i.e., encoder output) and the GP-modulated topic proportions without relying on an explicit evidence lower bound. The third term is the KL divergence regularizing the parameters of the approximate GPs: as derived in [24], it has a closed-form expression (Appendix B). A scaling coefficient is applied to balance its contribution with the other loss terms. The total loss is minimized via stochastic gradient descent with mini-batching [45]. This formulation allows LGTM to be trained in an end-to-end differentiable manner while incorporating covariates to modulate the latent space.

### 2.5. Hyperparameter Selection and Covariate Importance

The number of topics, i.e., the latent dimension *L*, is an important hyperparameter of a topic model. Existing methods based on held-out likelihood [46] or topic dissimilarity [47] often do not exhibit a clear optimum and tend to favor large values of *L*. This contrasts with typical microbiome studies where a small *L* is preferred to maintain biological interpretability [32, 33]. We therefore view *L* as controlling the resolution of the topic representation and encourage choosing it in accordance with the biological question and downstream interpretation. As a complementary practical heuristic, we use profile likelihood [48] to detect the “elbow” point in the curve of reconstruction loss versus *L*. We further assess topic robustness with respect to *L* using topic alignment across a range of latent dimensions. See Appendix C.1 and Appendix C.2 for details.

To quantify the contribution of covariates to the model, we use a variance-based sensitivity analysis [49, 50]. Specifically, for each latent dimension, we compute the total Sobol’ index of each covariate for the corresponding dimension of the multi-output latent function *g*(*x*) (Eq. (4)). A large value indicates that the covariate has a strong influence on the variance of that specific topic. Because Sobol’ indices are defined relative to the variance of each output, we further multiply each index by the variance of the corresponding latent dimension. This variance weighting places the indices on a common variance scale, such that values are directly comparable across both latent dimensions and covariates. In addition, we compute the aggregated Sobol’ index [51] to obtain an overall importance score per covariate, by averaging the topic-specific indices, weighted by the variance of each dimension’s output.

## 3. Experiments and Results

### 3.1. Datasets

We evaluated our model and demonstrated its application on three publicly available microbiome datasets with various characteristics. Each dataset comprises *P* subjects sampled at up to *T* timepoints, as summarized in Table 1. The Dhaka dataset contains metagenomic data from a cohort of *P* = 222 young children from Dhaka, Bangladesh. Taxonomic profiles were generated by MetaPhlAn2 [41]. The DIABIMMUNE dataset contains 16S rRNA amplicon sequencing data from a study of young children in Finland, Estonia, and Russia, focusing on the early gut microbiome and its associations with autoimmune development and aberrations. The HMP2 dataset contains metagenomic data from the Integrative Human Microbiome Project, which focuses on the gut microbiome in inflammatory bowel disease (IBD). See Appendix D for more details.

**Table 1:**
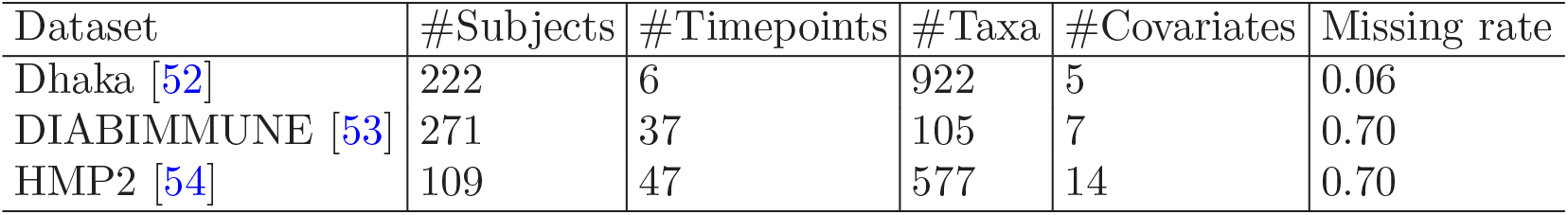
Summary of longitudinal datasets. Each dataset includes *P* subjects sampled at up to *T* timepoints. #Taxa and #Covariates denote the number of taxa and meta-data variables used in modeling. Missing rate is computed as 1 − *N/*(*P* × *T*), where *N* is the number of observed samples.

### 3.2. LGTM Provides Accurate Imputation and Forecasting of Longitudinal Microbial Profiles

We conducted imputation and forecasting experiments to evaluate LGTM’s ability to perform temporal interpolation and future prediction. These supervised learning tasks involve predicting relative abundances at missing or future timepoints given observed data and covariates. We focused on the Dhaka and DIABIMMUNE datasets from young children for these tasks, as the microbiome in early life undergoes substantial changes and provides clear temporal signals for imputation and forecasting. We compared our model against several methods ranging between naive and advanced techniques, including the recently proposed deep generative basis function Gaussian process (DGBFGP) [24]. Since both the observed *y* and the predicted 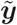 can be interpreted as probability vectors, we used cross-entropy as an evaluation metric, equivalent to the negative log-likelihood under a multinomial assumption. We reported the average test performance across multiple training-validation-test splits and multiple random seeds of the models. For imputation, we used sample-level splits (70/10/20); the training set includes samples from all subjects to model subject ID. For forecasting, we used subject-level splits (10% validation, 20% test); for validation/test subjects we input the first half of timepoints and evaluated on the second half. See Appendix E for more details.

Other studies [16, 18] apply transformations to the data for training and evaluation, such as the centered log-ratio (CLR) transform [55], which requires adding pseudo-counts to handle unobserved taxa (i.e. zeros) [56]. We found that models optimized for mean squared error (MSE) on log-transformed data performed poorly when their predictions were converted back to relative abundances (Appendix E.3). We therefore assessed predictive performance directly on the compositional data using likelihood, following the convention in topic modeling. This approach avoided potential biases introduced by log-transformation and aligned with the model’s probabilistic formulation. MSE for the compositional data was included in the results for reference.

Overall, our model outperformed the RNN-based BRITS and attention-based SAITS, which are widely used for time-series imputation, demonstrating the advantage of using GPs to capture covariate dependencies (Table 2). Compared to the state-of-the-art DGBFGP, our model provided a favorable trade-off by replacing the black-box MLP decoder with an interpretable topic model linear decoder. This design provides biological interpretability in the latent space, while incurring a minor decrease in predictive accuracy.

**Table 2:** Imputation and forecasting performance on two children datasets (Dhaka and DIABIMMUNE). Impute denotes predicting missing timepoints, and Forecast denotes predicting future timepoints given early observations. Both tasks are trained with observed microbiome profiles, but at prediction time the model uses covariates only. NLL (↓) is the negative log-likelihood, and MSE (↓) is reported in 10^−3^ for reference. Results are averaged over multiple train/validation/test splits and random seeds. Results for the best and second-best methods are bolded and underlined, respectively.

|  | Dhaka |  |  |  | DIABIMMUNE |  |  |  |
| --- | --- | --- | --- | --- | --- | --- | --- | --- |
|  | Impute |  | Forecast |  | Impute |  | Forecast |  |
|  | NLL | MSE | NLL | MSE | NLL | MSE | NLL | MSE |
| Mean | 3.34 | 0.215 | 3.90 | 0.190 | 2.92 | 1.32 | 2.89 | 0.90 |
| Mean (subjectwise) | 4.56 | 0.276 | 8.17 | 0.372 | 2.87 | 1.12 | 3.53 | 1.18 |
| Linear interpolation | 4.96 | 0.247 | 8.47 | 0.296 | 2.85 | <u>1.00</u> | 3.76 | 1.13 |
| BRITS [13] | 3.47 | 0.202 | 4.12 | 0.150 | 2.73 | 1.03 | 2.76 | 0.76 |
| SAITS [15] | 3.32 | 0.196 | 3.71 | <u>0.135</u> | 2.81 | 1.21 | 2.77 | 0.78 |
| DGBFGP [24] | **3.07** | **0.148** | **3.62** | **0.126** | **2.60** | **0.88** | **2.72** | **0.70** |
| LGTM | <u>3.11</u> | <u>0.150</u> | <u>3.68</u> | <u>0.135</u> | <u>2.69</u> | 1.01 | <u>2.75</u> | <u>0.73</u> |

### 3.3. LGTM Discovers Diverse and Stable Microbial Topics

We next evaluated the quality of the topics discovered by the model using all three datasets. Our approach was compared with topic modeling methods that explicitly yield topic-taxon distributions, including classical approaches used in microbiome studies (NMF and LDA). We also included VAE-LDA, a neural topic model that does not incorporate covariates. Cross-validation was performed to assess the goodness of fit using likelihood. To assess topic quality, we measured diversity and stability. For diversity, we used the Jensen–Shannon (JS) divergence and Bray–Curtis dissimilarity to compute pairwise similarities between topics. We further defined the distance between two topic matrices as the minimized average JS distance [57] between matched topics, solving an assignment problem [58]. The pairwise topic matrix distance was then computed to quantify the stability of the model across runs with different random seeds. This task can be viewed as reconstruction, which is less challenging than prediction, since *y* is available during testing. See Appendix F for more details.

LGTM outperformed other models in topic diversity over a wide range of *L* (Figure 2). This indicates that there was consistently lower redundancy between topics discovered by LGTM compared to other models. The higher LGTM topic diversity is likely related to the simplex-level KL alignment between encoder-derived and GP-modulated topic proportions, which encourages diverse topics while maintaining reconstruction quality. In line with this, all the methods achieved a similar likelihood, indicating comparable reconstruction quality (Table 3).

**Table 3:** Topic discovery performance on three datasets. This task is reconstruction: topic proportions are inferred from observed microbiome profiles, and NLL (↓) is the cross-validated reconstruction negative log-likelihood. JSD (↑) is the minimum pairwise Jensen–Shannon divergence, and BCD (↑) is the minimum pairwise Bray–Curtis dissimilarity, both measuring topic diversity (higher is better). TD (↓) is the maximum pairwise topic matrix distance, measuring stability across runs (lower is better). nnd denotes the NNDSVD initialization. Results are averaged over multiple random seeds. Results for the best and second-best methods in diversity and stability are bolded and underlined, respectively.

|  | Dhaka |  |  |  | DIABIMMUNE |  |  |  | HMP2 |  |  |  |
| --- | --- | --- | --- | --- | --- | --- | --- | --- | --- | --- | --- | --- |
|  | NLL | JSD | BCD | TD | NLL | JSD | BCD | TD | NLL | JSD | BCD | TD |
| NMF [26] | 2.74 | 0.747 | 0.843 | **0.028** | 2.46 | 0.758 | 0.846 | **0.021** | 3.03 | 0.478 | 0.610 | **0.028** |
| NMF (w/o nnd) | 2.74 | 0.761 | 0.850 | 0.166 | 2.47 | 0.621 | 0.722 | 0.384 | 3.03 | 0.515 | 0.641 | 0.529 |
| LDA [25] | 2.74 | 0.745 | 0.833 | 0.446 | 2.44 | 0.749 | 0.846 | 0.293 | 3.01 | 0.560 | 0.670 | 0.575 |
| VAE-LDA [27] | 2.77 | 0.764 | 0.846 | 0.489 | 2.45 | 0.666 | 0.797 | 0.344 | 3.06 | 0.648 | 0.777 | 0.614 |
| LGTM (w/o nnd) | 2.77 | <u>0.835</u> | **0.887** | 0.452 | 2.47 | **0.870** | **0.917** | 0.521 | 3.07 | **0.921** | **0.951** | 0.601 |
| LGTM | 2.74 | **0.840** | <u>0.877</u> | <u>0.124</u> | 2.45 | <u>0.796</u> | <u>0.871</u> | <u>0.142</u> | 3.03 | <u>0.854</u> | <u>0.904</u> | <u>0.081</u> |

**Figure 2:**
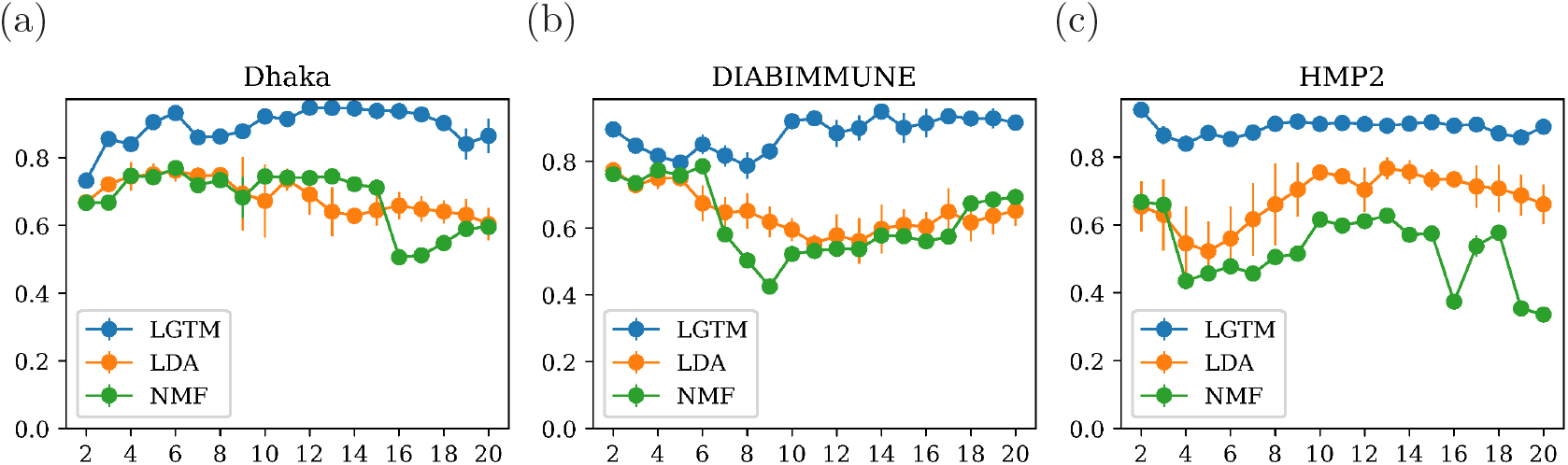
LGTM discovered diverse topics across a wide range of *L*. Topic diversity versus the number of topics on (a) Dhaka, (b) DIABIMMUNE and (c) HMP2 profiles. Diversity was measured as the minimum pairwise Jensen–Shannon divergence across topics. Points indicate mean across random seeds and error bars (vertical lines) indicate one standard deviation.

Stochasticity stemming from model initialization is a critical challenge for topic models, potentially leading to different topic matrices across runs, making their interpretation somewhat arbitrary. It is known that NMF becomes nearly deterministic when initialized with NNDSVD to improve approximation. We adapted this technique to initialize the LGTM decoder and observed a substantial improvement in stability. This demonstrates that consistent initialization is a crucial step toward achieving reproducible results in deep generative topic models.

### 3.4. LGTM Identifies Interpretable Microbial Co-abundance Patterns

Having shown quantitatively that our approach can effectively capture temporal dynamics while discovering diverse topics with improved stability across runs, we now qualitatively demonstrate the interpretability and utility of our model.

We first utilized metagenomes from young children in Dhaka, Bangladesh to model microbial succession and its associations with early feeding (Figure 3). LGTM learned sparse topics each characterized by a few distinct taxa. This sparsity is beneficial for interpretation, as it allows us to describe the data using a small number of key taxa that tend to either co-occur or have distinct abundance patterns. Unsurprisingly, child’s age accounted for the largest variation in topic proportions, indicating rapid gut microbiome development during the first two years of life. Breastfeeding had a relatively large influence on the composition of the microbiome, especially on the most prominent Topic 1, which was dominated by members of the genus *Bifidobacterium*.

**Figure 3:**
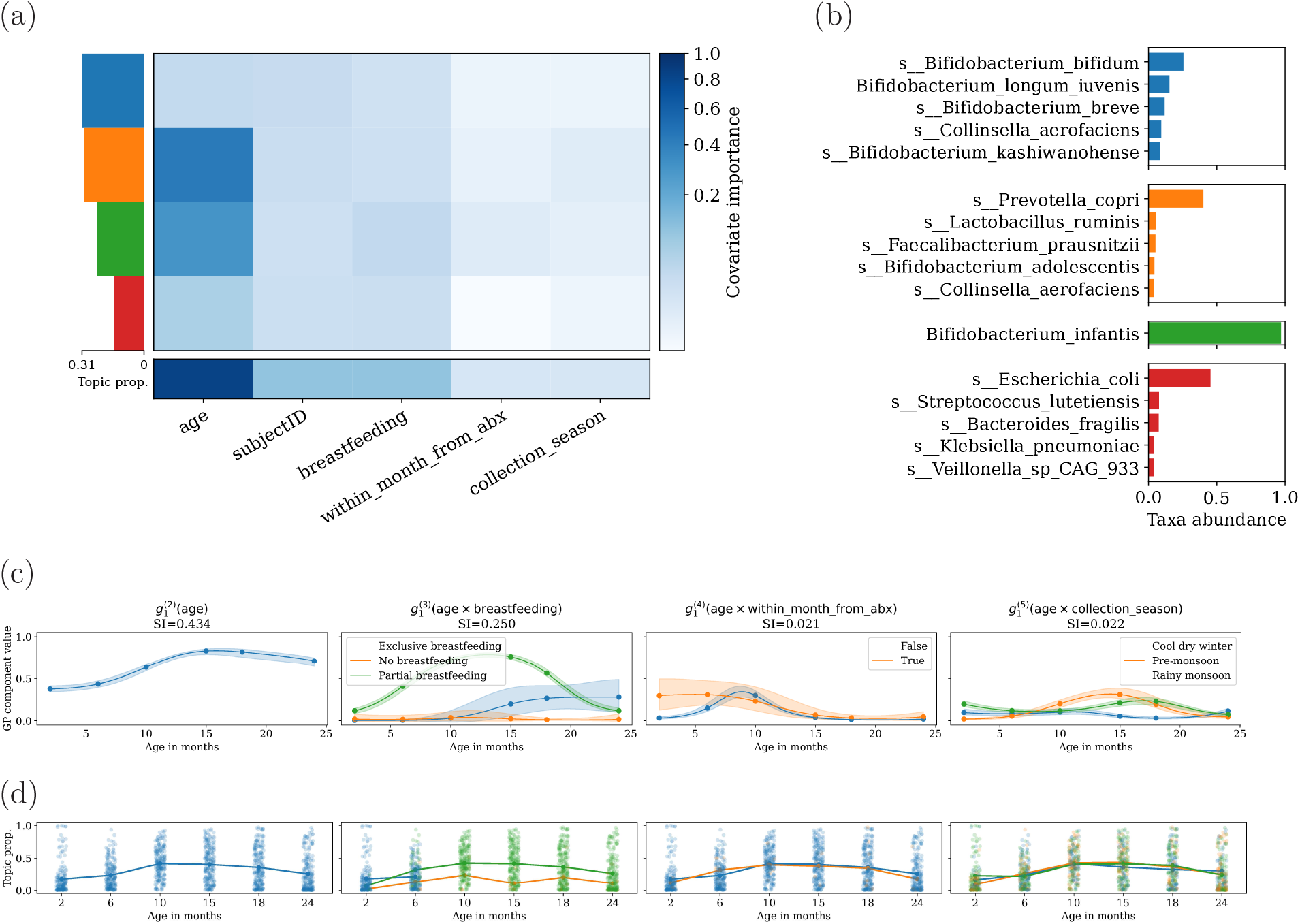
LGTM captured longitudinal trends and topics associated with early feeding in young children from Dhaka, Bangladesh. (a) Heatmap of variance-weighted Sobol’ indices quantifying the contribution of each covariate (columns) to the variation of each topic’s proportion (rows, *L* = 4). The bottom row shows the aggregated Sobol’ indices for each covariate across all topics. Topics are sorted by their mean proportion across all samples, indicated by the horizontal bars on the left. (b) Taxonomic composition of the learned topics. For each topic, the bar chart displays the top five most abundant taxa whose proportions exceed 0.01. (c) Latent functions for Topic 1. The learned latent function is decomposed into components. Each subplot shows the effect of a single covariate or an interaction term on the topic’s proportion over age in months. The subject effect is not shown, as it reflects individual differences rather than age-related variation. Shaded areas represent one standard deviation of uncertainty. The Sobol’ index (SI) is displayed above each subplot. For interaction components, the SI of the categorical variable is shown. (d) Strip plots of topic proportions over timepoints. Topic proportions are computed via the encoder. Each point represents a sample, colored according to the legends in the upper subplots. Lines connect the mean topic proportion at each timepoint for each category, illustrating the trends that are modeled by the latent functions in the subplots above.

By further visualizing the latent functions for Topic 1, we observed that breastfeeding status exhibited contrasting temporal interactions with Topic 1, whereas recent antibiotic courses and collection season showed milder effects (Figure 3c, d). These patterns were successfully captured by the corresponding components of the latent function. Topic 1 was more prevalent when the infants were introduced to solid foods, in line with previous findings that *Bifidobacterium longum* subsp. *iuvenis* thrived during the transition from exclusive to partial breastfeeding [52, 59]. During the same period, the abundance of *Bifidobacterium longum* subsp. *infantis* decreased while *Prevotella copri* started to increase, as captured by Topic 2 (Figure G1). Altogether, LGTM captured complex bacterial co-abundance patterns and their associations with age and other covariates while at the same time providing easy-to-access visual representations of these trends.

We next applied our model to the 16S rRNA amplicon sequence data from the integrated DIABIMMUNE cohort of young children in Finland, Estonia and Russian Karelia. In these data, the country of birth accounted for a large portion of the variation for several topics (Figure 4). The genus *Bacteroides* was more prevalent and abundant in Finnish and Estonian children, as reported previously [60]. On the other hand, LGTM further identified the lack of early Bacteroides in children born by C-section (Figure 4c), which is a well-established effect from numerous studies [61, 62]. LGTM further captured major trends in the genus *Bifidobacterium*; Russians had more bifidobacteria in the early months and, on the other hand, breastfeeding tended to increase *Bifidobacterium* levels, as breast milk provides an energy source primarily for these bacteria.

**Figure 4:**
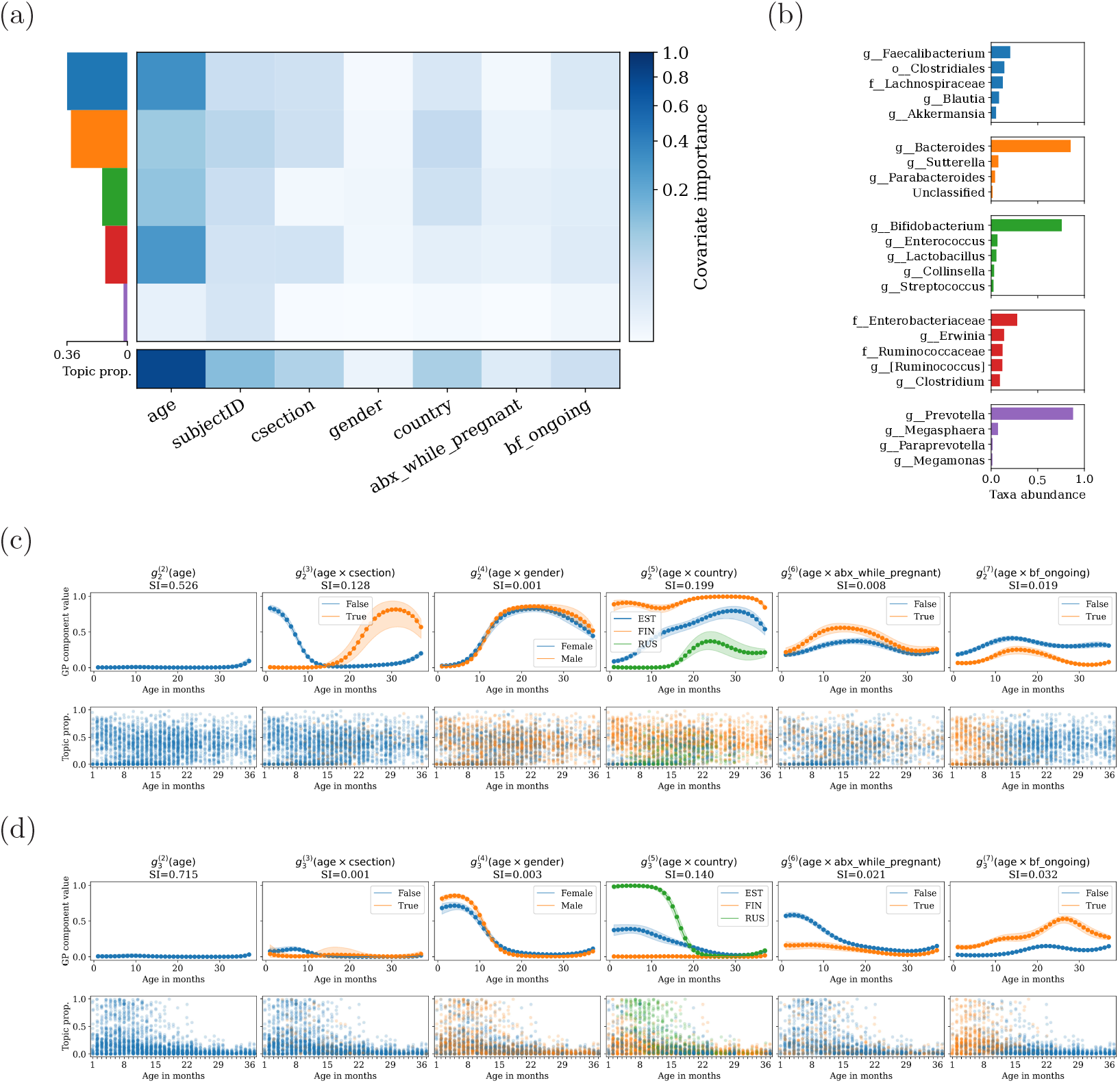
LGTM facilitated deconvolution of microbiome shifts related to complex early life events in the DIABIMMUNE cohort. (a) Heatmap of variance-weighted Sobol’ indices quantifying the contribution of each covariate (columns) to the variation of each topic’s proportion (rows, *L* = 5). The bottom row shows the aggregated Sobol’ indices across all topics. Topics are sorted by their mean proportion across all samples, indicated by the horizontal bars on the left. (b) Taxonomic composition of the learned topics. For each topic, the bar chart displays the top five most abundant taxa whose proportions exceed 0.01. (c) Latent functions and topic proportions for Topic 2. Upper panels: each subplot shows the effect of a single covariate or an interaction term on the topic’s proportion over age in months. The subject effect is not shown, as it reflects individual differences rather than age-related variation. Shaded areas represent one standard deviation of uncertainty. The Sobol’ index (SI) is displayed above each subplot. For interaction components, the SI of the categorical variable is shown. Lower panels: strip plots of topic proportions over timepoints. Topic proportions are computed via the encoder. Each point represents a sample, colored according to the legends in the upper panels. (d) Latent functions and topic proportions for Topic 3, with the same visualization scheme as in (c). Legend: csection, caesarean section; EST, Estonia; FIN, Finland; RUS, Russia; abx while pregnant, antibiotic use during pregnancy; bf ongoing, breastfeeding.

We further applied our model to the metagenomic data from the Human Micro-biome Project (phase 2) [54]. Unlike gut microbiome profiles from young children, time accounted for a relatively small proportion of variation in these IBD-associated microbiome profiles, while inter-subject variation accounted for the largest portion of the variance (Figure 5). This was mirrored with strong heterogeneity of the gut microbiome profiles across subjects. Subjects were classified into three groups based on their disease diagnosis: Crohn’s disease (CD), ulcerative colitis (UC), and controls without inflammatory bowel disease (non-IBD). While the variance contribution of disease diagnosis was moderate, that of dysbiosis state (as reported in [54]) was larger, indicating that LGTM distinguished dysbiotic and non-dysbiotic microbiome profiles. Microbiome features distinctive to dysbiotic microbiome profiles were distributed across Topics 1 and 5. Topic 1 was characterized by species often prevalent and abundant in healthy individuals. For example, *Bacteroides uniformis* and *Alistipes putredinis* are often depleted in IBD and their protective roles against IBD have been investigated [63, 64]. Topic 5 captured species increased in dysbiosis, such as *Escherichia coli* and *Ruminococcus gnavus*. There was a weaker association to IBD diagnosis with Topics 1, 2 and 3. Topic 2 was enriched in both UC and CD whereas enrichment of Topic 3 was characteristic of UC. Topics 2 and 3 were characterized by *Bacteroides vulgatus* and *Faecalibacterium prausnitzii*, respectively, both of which have been extensively studied in IBD. Overall, while the topics showed large variance across subjects, they were also associated with other covariates. For example, Topic 1 was associated with probiotic intake, and consumption of drinks with artificial sweeteners, dairy and red meat, whereas Topic 6 was associated with having whole grains in diet. Altogether, this demonstrates LGTM’s ability to capture disease-relevant microbiome shifts while jointly modeling diet and other measured metadata.

**Figure 5:**
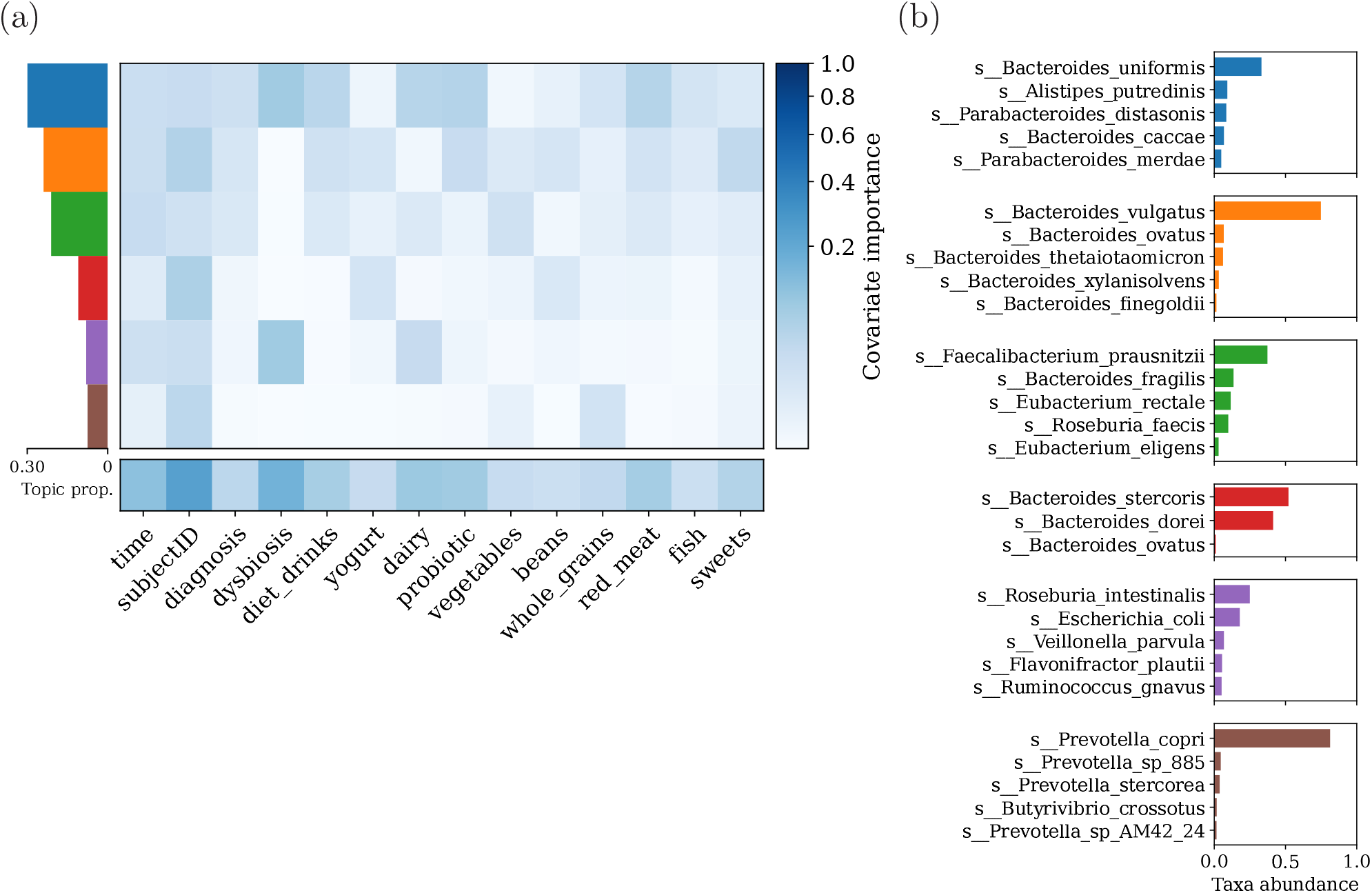
LGTM topics were associated with patient diagnosis, microbial dysbiosis and dietary factors in HMP2 data. (a) Heatmap of variance-weighted Sobol’ indices quantifying the contribution of each covariate (columns) to the variation of each topic’s proportion (rows, *L* = 6). The bottom row shows the aggregated Sobol’ indices across all topics. Topics are sorted by their mean proportion across all samples, indicated by the horizontal bars on the left. (b) Taxonomic composition of the learned topics. For each topic, the bar chart displays the top five most abundant taxa whose proportions exceed 0.01.

## 4. Discussion

We present LGTM, a probabilistic learning framework for modeling longitudinal microbiome data with external covariates, which combines the flexibility of GPs with the interpretability of topic models. LGTM represents the microbiome using interpretable topics, each defined by a small set of co-varying microbial features, typically species. Importantly, topics are learned jointly with external covariates allowing them to capture correlations with host and environmental factors in a data-driven manner. The unified probabilistic framework enables direct comparisons of microbial trajectories across individuals and conditions while accounting for uncertainty, compositionality, and irregular sampling. In contrast to methods that analyze taxa independently, the topic-based structure in LGTM facilitates biological interpretation by linking microbial topics to measured host and environmental covariates. Using multiple datasets, we demonstrated that this allows identifying distinct microbial topics with coherent temporal behavior and association with relevant host covariates, including disease state and dietary changes. Together, these results demonstrate that LGTM provides an interpretable and flexible framework for studying longitudinal gut microbiome dynamics in the presence of complex covariates.

Our primary technical contribution is a deep generative framework that, to our knowledge, is the first to integrate basis function approximation for GPs with topic modeling in an autoencoder architecture. A central feature is interpretable dimensionality reduction. Samples are represented as mixtures of latent factors, which decompose co-varying taxa into structured co-abundance patterns. Additive GPs summarize associations with continuous and categorical covariates in the latent space, while a simplex-constrained linear decoder enhances interpretability by directly linking the latent factors to the original high-dimensional features. The GP components decompose covariate contributions, enabling clearer biological interpretation of covariate associations in longitudinal data. Together, these provide two complementary decompositions of longitudinal microbiome variation, capturing co-abundance-defined microbial topics and covariate contributions to support multi-faceted interpretability.

The inferred topics represent statistical co-abundance patterns rather than direct evidence of ecological guilds, functional groups, or pairwise microbial interactions. When the taxa in a topic are biologically coherent and the topic is associated with relevant covariates, it can suggest a hypothesis about microbial community structure, supported by external biological knowledge or downstream validation. Mechanistic ecological models, such as gLV models, address a different level of analysis by parameterizing growth and interaction coefficients. The topic proportions inferred by LGTM could therefore serve as lower-dimensional inputs for other analyses that combine topic modeling with explicit ecological interaction models.

Several statistical assumptions define the scope of the current model. LGTM is trained on taxonomic profiles represented as relative abundances, and the cross-entropy reconstruction term can be viewed as a multinomial-style working likelihood on the simplex. This preserves compositional structure and avoids CLR/log-ratio preprocessing that requires zero replacement or pseudocounts, but it does not explicitly model sample-specific read depth, overdispersion, or abundance-level zeros. Dirichlet– multinomial, zero-aware logistic-normal, or other count-aware likelihoods are natural extensions when raw count-generative modeling is required. For temporal modeling, the squared exponential kernel is appropriate for gradual longitudinal trends, and perturbation-related metadata can enter the model through covariate-time interactions, but abrupt unrecorded or poorly sampled changepoints may be smoothed over. Other stationary kernels compatible with Hilbert-space basis-function approximations, such as Matérn kernels, could relax this smoothness assumption. When precise perturbation timings are available, event-aligned covariates, such as time since antibiotic exposure or time since dietary change, or input-warped kernels based on stationary kernels would be natural extensions for modeling trajectories relative to the perturbation rather than only over absolute time [65]. Finally, the additive GP structure and Sobol’ indices summarize covariate associations rather than causal effects, so results for correlated metadata variables should be interpreted with care. Practical steps, such as applying general-purpose feature selection methods to select a representative subset of covariates, can be combined with the same modeling framework when correlated metadata are central to the analysis. In future implementations, feature selection could also be incorporated during model fitting through sparsity-inducing penalties or shrinkage priors on covariate-specific GP components.

Selecting the latent dimensionality (i.e., the number of topics) remains an open practical question [37]. We view *L* primarily as a resolution parameter: smaller values provide a compact summary of major co-abundance structure, whereas larger values can retain more specific or lower-variance patterns when they are relevant for the analysis. The choice of *L* should therefore be guided by the biological question, downstream interpretation, and sensitivity analysis. In addition, topic models may have identifiability issues because random initialization can affect the detailed composition of topics. In our implementation, NNDSVD initialization provides a useful anchor for the decoder by orienting topics toward dominant taxon-level patterns, improving robustness of the major topics without eliminating the need for biological interpretation.

The neural architecture is extensible and supports several directions for future work. For example, while we consider covariates in the metadata, one could also include response variables (such as clinical outcomes), which could then be predicted using the microbiome profiles or their latent representations. Other data modalities, such as metabolomics, could also be incorporated to enable more comprehensive modeling. At the same time, potential overfitting is mitigated by the low latent dimension and the simplex-constrained linear decoder, which constrains effective model capacity while preserving interpretability. The held-out predictive performance supports this constrained-capacity interpretation. Phylogenetic information could further strengthen biological interpretation, for example through a tree-structured prior or regularization on the topic-taxon matrix, inspired by Dirichlet-tree multinomial models [66]. Functional information could similarly be incorporated by jointly modeling functional profiles from tools such as HUMAnN, MG-RAST, or PICRUSt2 as an auxiliary output [67, 68, 69]. A limitation is that, due to the heterogeneity of microbiome and metadata across cohorts, our model currently focuses on analyzing data from one cohort at a time. Generalization across cohorts remains a significant challenge in microbiome studies, and transferable topics would require a shared taxonomic feature space, comparable metadata, and explicit validation that matched topics preserve similar biological interpretation across cohorts. As large-scale integrative microbiome studies expand, the scalability of our approach will become increasingly valuable. Overall, LGTM provides a scalable and interpretable foundation for analyzing longitudinal microbiome data, identifying biological patterns, and generating hypotheses for future experimental validation.

## Acknowledgements

We would like to acknowledge the computational resources provided by the Aalto Science-IT. We acknowledge CSC for awarding this project access to the LUMI super-computer, owned by the EuroHPC Joint Undertaking, hosted by CSC (Finland) and the LUMI consortium. ChatGPT (OpenAI, GPT-5.2) was used for language editing and clarity improvement.

## Code and data availability

The LGTM source code is available at https://github.com/yuanx749/lgtm. An archived version of the code is available on Zenodo at https://doi.org/10.5281/zenodo.20588704. Data from the DIABIMMUNE and Dhaka studies are available at NCBI Sequence Read Archive under Bio-Project IDs PRJNA497734 and PRJNA806984, respectively. DIABIMMUNE data can also be accessed through the DIABIMMUNE microbiome website at https://diabimmune.broadinstitute.org/. Microbiome profiles of the HMP2 study are publicly available via the IBDMDB at https://ibdmdb.org/.

## Appendix A. Basis function GP for continuous, categorical and interaction kernels

We follow earlier work to formulate basis function approximation for GPs [39, 40, 24]. We use the SE kernel to model continuous covariates, which is approximated as:

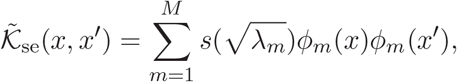

where *s*(·) is the spectral density, *λ*_*m*_ and *φ*_*m*_(*x*) are the *m*-th eigenvalue and eigen-function of the Laplace operator. The corresponding GP has the following parametric form:

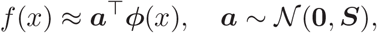

where *φ*(*x*) = [*φ*_1_(*x*), …, *φ*_*M*_ (*x*)]^⊤^ and 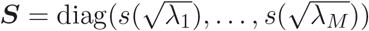. Note that the approximation depends on the kernel hyperparameters. The SE kernel is defined as:

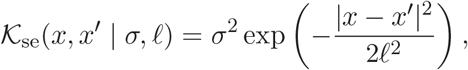

where *σ*^2^ is the variance determining the overall variation of the function and *ℓ* is the length scale determining how quickly the function values change with distance. Its corresponding spectral density is

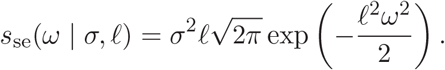

We model categorical covariates using a categorical kernel, which is constructed from the eigendecomposition of the kernel matrix. Let *x* ∈ *{*1, …, *C}* be a categorical covariate with *C* categories, and let *K*_ca_(*x, x*^′^) ℝ^*C*×*C*^ denote the kernel function. A symmetric kernel matrix admits an eigendecomposition and the kernel function can be written as:

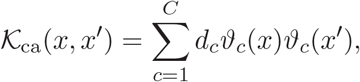

where *d*_*c*_ is the *c*-th eigenvalue and *ϑ*_*c*_(*x*) is the *x*-th entry of the *c*-th eigenvector of the corresponding kernel matrix. The corresponding GP then has an exact parametric form:

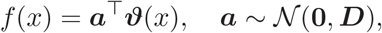

where *ϑ*(*x*) = [*ϑ*_1_(*x*), …, *ϑ*_*C*_(*x*)]^⊤^ and *D* = diag(*d*_1_, …, *d*_*C*_). In our implementation, we specifically use the diagonal kernel for categorical variables:

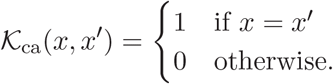

To model the joint effects of continuous and categorical covariates, we compute the product of the approximation for the SE kernel and the decomposition of the categorical kernel:

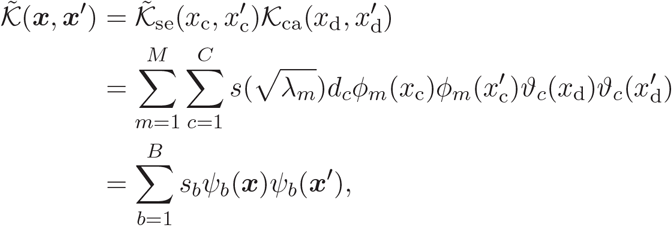

where 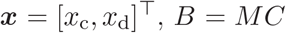, and each *s*_*b*_ and *ψ*_*b*_ represents one of the products in 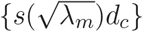 and {*φ*_*m*_(*x*_c_)*ϑ*_*c*_(*x*_d_)}, respectively. Note that 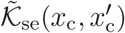 depends only on a single continuous variable and 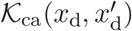 depends only on a single categorical variable. The corresponding GP with this interaction kernel has the following parametric form:

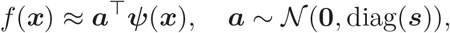

where *ψ*(*x*) = [*ψ*_1_(*x*), …, *ψ*_*B*_(*x*)]^⊤^ and *s* = [*s*_1_, …, *s*_*B*_]^⊤^.

## Appendix B. KL divergence for the basis function GP approximation

The second regularization term in our training objective in Eq. (5), KL_GP_, is derived in earlier work [24], which we describe here. Let *A, σ, ℓ* denote the collection of parameters *A*^(*r*)^, *σ*^(*r*)^, *ℓ*^(*r*)^ for all components *r* = 1, …, *R*. These parameters have the following prior and variational distributions:

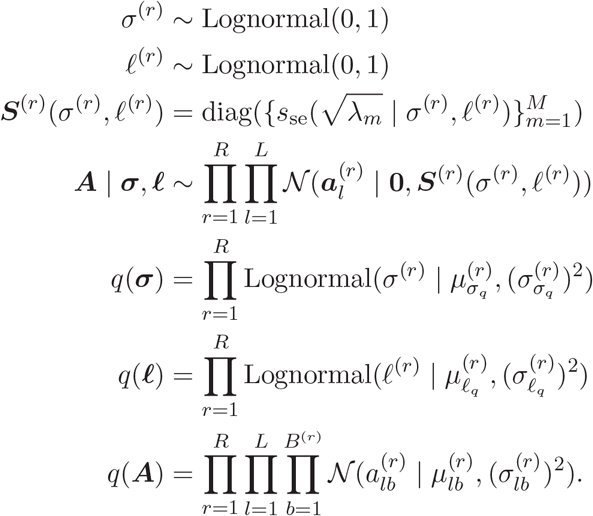

During inference, we sample *σ* and *ℓ* from the corresponding distributions. These values are then substituted into the spectral density function to obtain an estimate, denoted as *ŝ*. Using a single Monte Carlo sample and the mean-field approximation, the KL divergence is computed as:

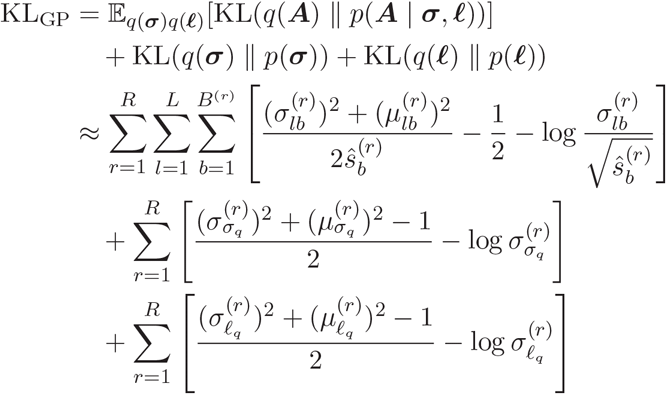

## Appendix C. Additional details of hyperparameters

### C.1. Latent dimension selection via profile likelihood

Let *e*_*L*_ be an evaluation metric, such as the average reconstruction loss from cross-validation, given latent dimension *L*. The heuristic assumes that for *L < L*^∗^, where *L*^∗^ is the (unknown) correct number of topics, the metric *e*_*L*_ will decrease rapidly, whereas for *L > L*^∗^, the marginal gains become smaller as the model is sufficiently complex. This method partitions the evaluation values {*e*_*L*_*}* into two groups at a changepoint *L* and assumes a simple model: *e*_*k*_ ∼ *N* (*µ*_1_, *σ*^2^) if *k < L* and *e*_*k*_ ∼ *N* (*µ*_2_, *σ*^2^) if *k* ≥ *L*. Maximum likelihood estimates are computed for the parameters, and the profile log-likelihood *ℓ*(*L*) is obtained:

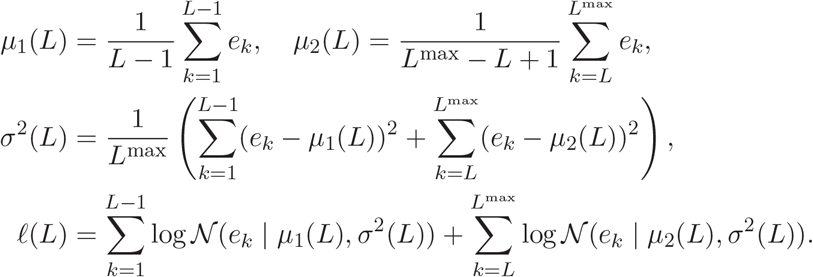

We use 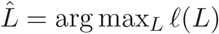 as a suggested topic number. For computational efficiency, this can also be applied to the reconstruction loss on the whole data, varying *L* while holding other hyperparameters at their default values.

### C.2. Sensitivity analysis of latent dimensionality

To assess how the inferred microbial structures change with the number of topics, we aligned LGTM topic models trained with *L* = 2, …, 20 using topic alignment [37]. We used the optimal transport alignment option, which aligns topics across adjacent values of *L* using topic-taxon compositions and the relative masses of topics. Figure C1 provides a visual summary of whether dominant topics persist across resolutions or split into more specific components as *L* increases.

### C.3. Other hyperparameters

We used *M* = 5 basis functions for the SE kernel approximation. The coefficient for the KL term of GP was set to 0.001. We set the default learning rate to 0.05, batch size to 64, and hidden dimension of MLP to 64.

## Appendix D. Additional details of datasets and preprocessing

### D.1. Dhaka [52]

This dataset consists of metagenomic shotgun sequencing data from fecal samples of Bangladeshi children in the Microbiota and Health Study, sampled during the first two years of life. Samples collected during diarrheal episodes were excluded from our analysis. The covariates used include age, subject ID, breastfeeding status, recent antibiotic use (within one month) and collection season.

### D.2. DIABIMMUNE [53]

This dataset consists of 16S rRNA gene amplicon sequencing data from the DIABIMMUNE study. DIABIMMUNE observed Finnish, Estonian and Russian children for three years from birth by monthly stool sampling. For our analysis, abundances were merged to the genus level, and any unclassified taxa were grouped into a single taxon. The covariates used include age, subject ID, mode of delivery, gender, country of birth, antibiotic courses during pregnancy, and breastfeeding status.

### D.3. HMP2 [54]

This dataset is from the Inflammatory Bowel Disease Multi’omics Database (IBD-MDB), part of the Integrative Human Microbiome Project, a study tracking partici-pants with and without IBD for one year. We used the metagenomic shotgun sequencing data from stool samples. The week number of sample collection since recruitment was used as the timepoint. The dataset features a rich set of metadata, including subject ID, IBD diagnosis, dysbiosis, and dietary information. Dietary recalls were binarized according to whether a food item was consumed in the past week.

## Appendix E. Additional details of prediction tasks

### E.1. Methods for comparison

#### E.1.1. Mean

The mean imputation method predicts missing values by calculating the mean of all observed values for each feature. We also compare against a subject-wise mean imputation, which calculates the mean of observed values within each subject.

**Figure C1:**
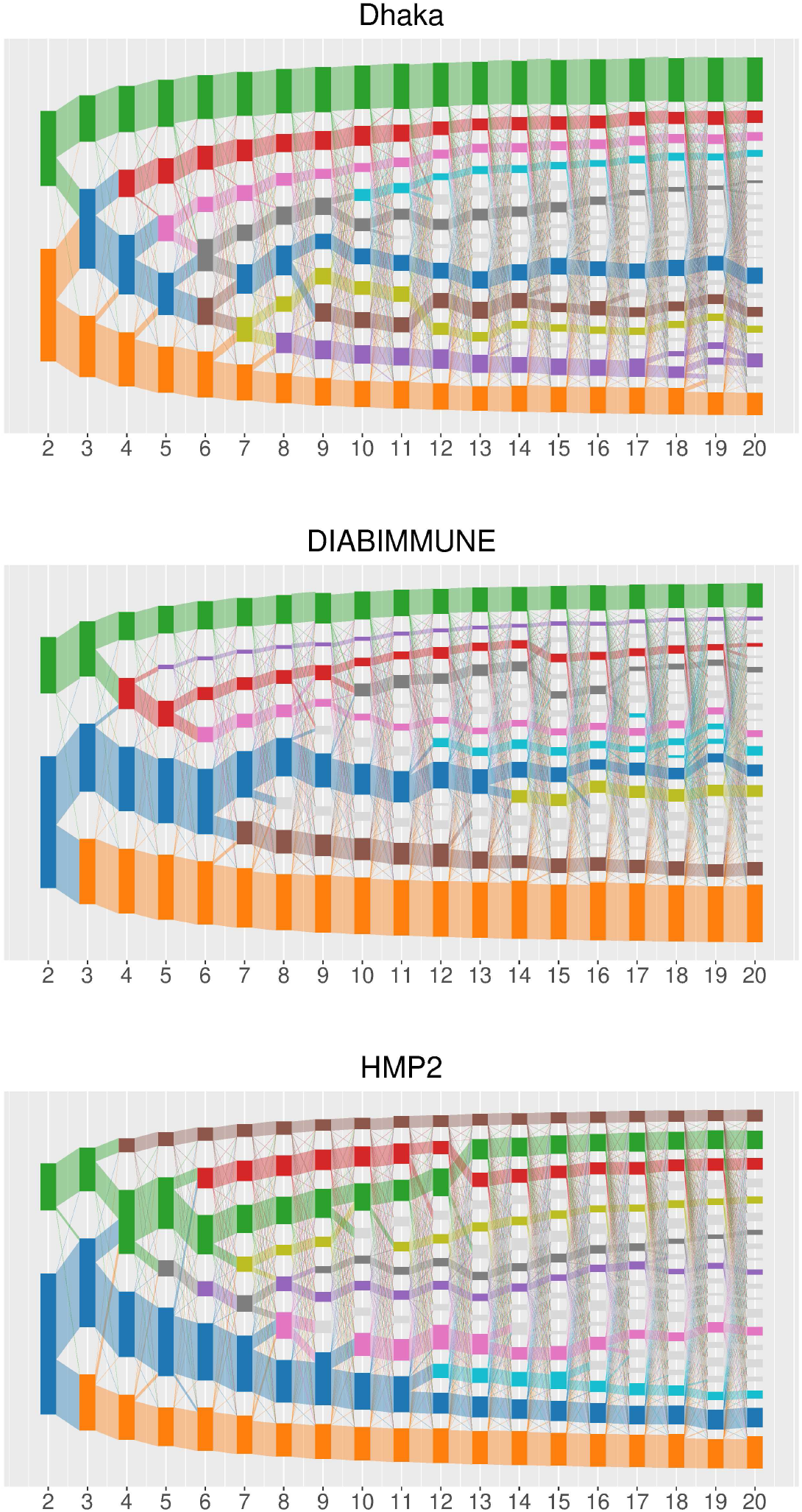
Robustness of the inferred topics with respect to the number of topics. Each panel shows LGTM models trained with *L* = 2, …, 20 topics for one dataset. Each rectangle represents one topic, and its height is proportional to the mean topic proportion across samples. Ribbons connect topics in adjacent values of *L*; ribbon widths indicate the optimal transport alignment mass between topic-taxon profiles. Colors are anchored at the topics from the main analyses. Additional high-mass paths are shown using the remaining colors, whereas minor paths are shown in grey. The alignment demonstrates that dominant microbial structures are robust across a range of *L*, while larger values of *L* mainly refine broader topics into smaller components.

#### E.1.2. Linear interpolation

This method predicts missing values by linearly interpolating between the nearest observed values before and after the missing timepoint for each feature within each subject. Missing values at the beginning or end of a series are backfilled or forward-filled with the nearest observed value.

#### E.1.3. BRITS [13]

BRITS is an RNN-based model that learns to impute missing values within a bidirectional recurrent dynamical system.

#### E.1.4. SAITS [15]

SAITS is a Transformer-based model that imputes missing values using a weighted combination of two diagonally-masked self-attention blocks. We used a configuration of two layers with four attention heads each.

#### E.1.5. DGBFGP [24]

DGBFGP is a deep generative model utilizing a Bayesian basis function approximation for its GP that eliminates the need for amortized variational inference. We used a MLP with one hidden layer for the decoder.

### E.2. Experimental details

For imputation, we split the data into training, validation, and test sets by samples with a ratio of 70%, 10%, and 20%, respectively. We performed five random splits such that each sample appeared in the test set of exactly one split, ensuring that all samples were evaluated. The training set contains samples from all subjects, which is necessary when modeling subject ID as a covariate, as the model must see each subject during training. For forecasting, we split subjects into validation (10%) and test (20%) sets. For subjects in the training set, all their timepoints were used; for subjects in the validation and test sets, only the first half of their timepoints were provided as input for training, while the second half was used for evaluation. We performed five random splits such that each subject appeared in the test set of one split.

Continuous covariates were standardized to have zero mean and unit variance for DGBFGP and our model. We implemented and trained all deep learning models in PyTorch [70]. The implementations in PyPOTS [71] were used for BRITS and SAITS. Cross-entropy loss was used for all models. Adam algorithm [45] was used for optimization. For each model, hyperparameters (e.g., learning rate, batch size, hidden dimension) were tuned using grid search guided by performance on the validation set. For each split, we used five random seeds for model initialization. We ran parallel hyperparameter optimization with Optuna [72], using multi-processing on a computing cluster to accelerate the search. We used a single thread for each process to ensure reproducibility. All models were trained for a maximum of 100 epochs.

### E.3. Experiment with CLR transform

The inputs were CLR-transformed relative abundances, replacing zeros with minimum non-zero value multiplied by 0.65 and multiplicatively adjusting the other values to ensure the compositions still add up to one. Models were trained with MSE loss, and hyperparameters were tuned based on validation MSE in the transformed space. The inverse CLR transform (i.e., softmax) was applied to the model outputs to obtain predictions in the original compositional space. As shown in Table E1, the likelihood is worse than training directly in the compositional space (Table 2), validating our methodological choice.

**Table E1:**
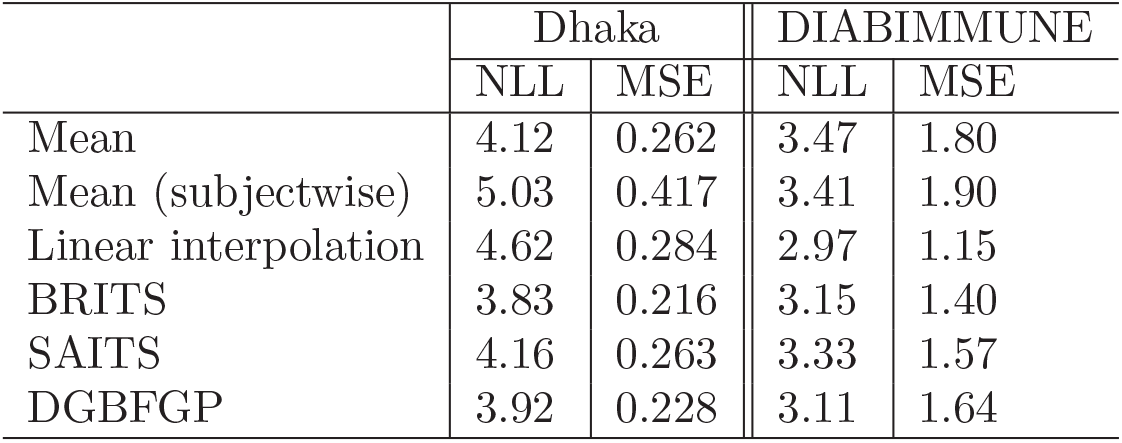
Imputation with CLR transform. NLL (↓): negative log-likelihood. MSE (↓): ×10^−3^.

## Appendix F. Additional details of topic discovery task

### F.1. Methods for comparison

#### F.1.1. NMF [26]

NMF finds two non-negative matrices whose product approximates the relative abundance matrix. Generalized KL divergence was used as the objective function to be minimized. The model was trained until convergence. The resulting matrices were normalized by dividing each row by its sum to obtain sample-topic and topic-taxon matrices.

#### F.1.2. LDA [25]

LDA is a generative probabilistic model using Dirichlet priors for sample-topic and topic-taxon distributions. It uses the variational Bayes method for parameter estimation. We trained the model for 100 epochs.

#### F.1.3. VAE-LDA [27]

VAE-LDA applies autoencoding variational Bayes to LDA. It uses Laplace approximation to approximate the Dirichlet prior for the sample-topic distributions. We used the same architecture for the encoder and decoder as in our model.

### F.2. Metrics

#### F.2.1. Likelihood

We assume a multinomial distribution for the abundance data with the same total count *K* for all samples. Let *k* = *Ky* be the observed counts and *ŷ* be the estimated relative abundances. The log-likelihood of the model is computed as:

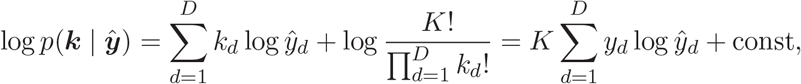

where *k*_*d*_, *y*_*d*_, and *ŷ*_*d*_ are the *d*-th elements of *k, y*, and *ŷ*, respectively. This is equivalent to the cross-entropy between the observed and estimated relative abundances up to a constant.

#### F.2.2. Jensen–Shannon divergence

The JS divergence is a symmetric version of KL divergence that measures the similarity between two probability distributions. For two topics *β*_*i*_ and *β*_*j*_, it is defined as:

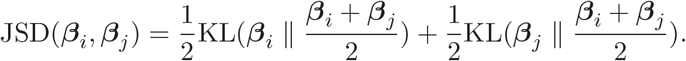

We used the base 2 logarithm, so it ranges from 0 to 1.

#### F.2.3. Bray–Curtis dissimilarity

Bray–Curtis dissimilarity is a measure of dissimilarity used in ecology to measure the difference in taxa composition between two samples. For two topics *β*_*i*_ and *β*_*j*_, it is defined as:

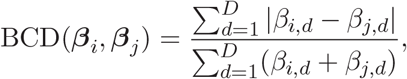

where *β*_*i,d*_ and *β*_*j,d*_ are the *d*-th elements of *β*_*i*_ and *β*_*j*_, respectively.

#### F.2.4. Topic matrix distance

We define the distance between two topic matrices *B*^(*r*)^ and *B*^(*s*)^ from runs *r* and *s* as the minimized average distance between matched topics. Let *π* be a permutation of {1, …, *L}* representing the bipartite matching between topics in the two matrices. The topic matrix distance is computed as:

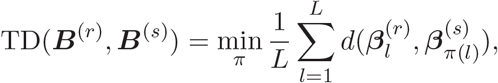

where *d*(·,·) is a distance metric. We used the JS distance (the square root of the JS divergence), which is a distance metric that satisfies the triangle inequality.

### F.3. Experimental details

We performed 5-fold cross-validation on samples to assess the held-out likelihood, using the same splitting strategy as in the imputation task. For the held-out likelihood calculation, the topic matrix (decoder) was held fixed, and the topic proportions for the test samples were optimized. To evaluate topic quality, we trained the models on all available data using five different random seeds. For NMF and LDA, implementations from scikit-learn [73] were used. We quantified the diversity of topics discovered using pairwise dissimilarity, with minimum pairwise dissimilarity to measure the worst-case redundancy and average pairwise dissimilarity to measure overall diversity. Our model consistently outperforms baselines over a wide range of topic numbers using different metrics (JS divergence and Bray–Curtis dissimilarity).

### F.4. Computational efficiency

Despite integrating sophisticated modules, our approach remains computationally efficient due to the basis function approximation for GPs. We further evaluated scalability using synthetic relative-abundance datasets by varying one dimension at a time while holding the others fixed. Runtime was measured as the total training time for 100 epochs on a single CPU thread, with *L* = 5 topics. Experiments were run on CPU with AMD EPYC 7763 processors. The benchmark varied the number of samples, taxa, subjects, and GP components. Runtime increased approximately linearly with the number of samples (log-log slope 1.02), showed only modest dependence on the number of taxa and subjects in the tested range (slopes 0.09 and 0.04), and increased with the number of GP components (slope 0.64; Figure F1). The sample-size dependence reflects the number of mini-batch updates per epoch and can be adjusted in practice by increasing the batch size. These results indicate that the main practical limit of the current implementation is the number and size of covariate components, since each component has its own basis-function expansion.

## Appendix G. Additional visualizations

### G.1. Dhaka dataset

**Figure F1:**
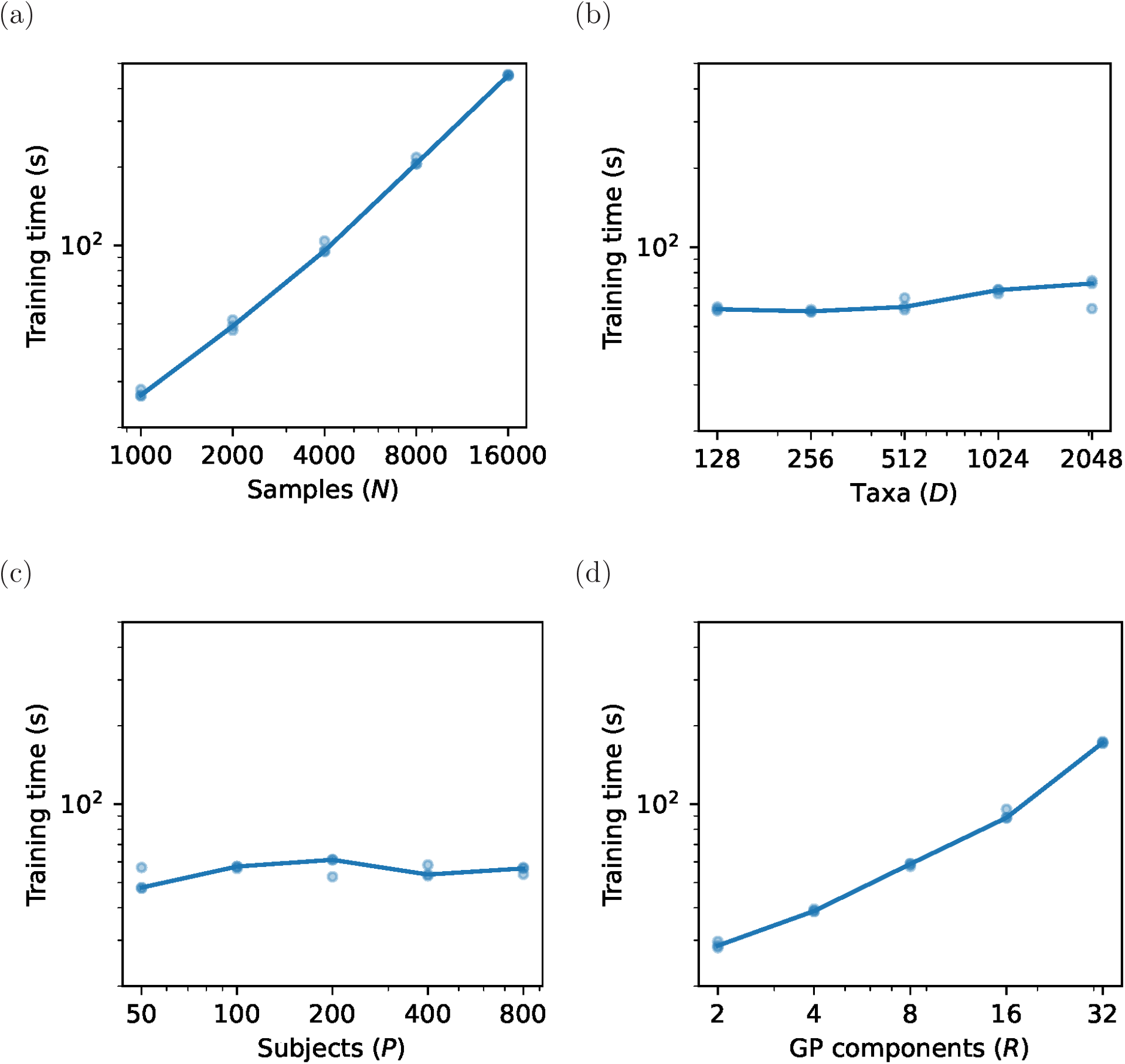
Scalability benchmark on synthetic data. Points show independent runs and lines show medians. One dimension was varied at a time while the other settings were fixed. Training time was measured for 100 epochs on a single CPU thread.

**Figure G1:**
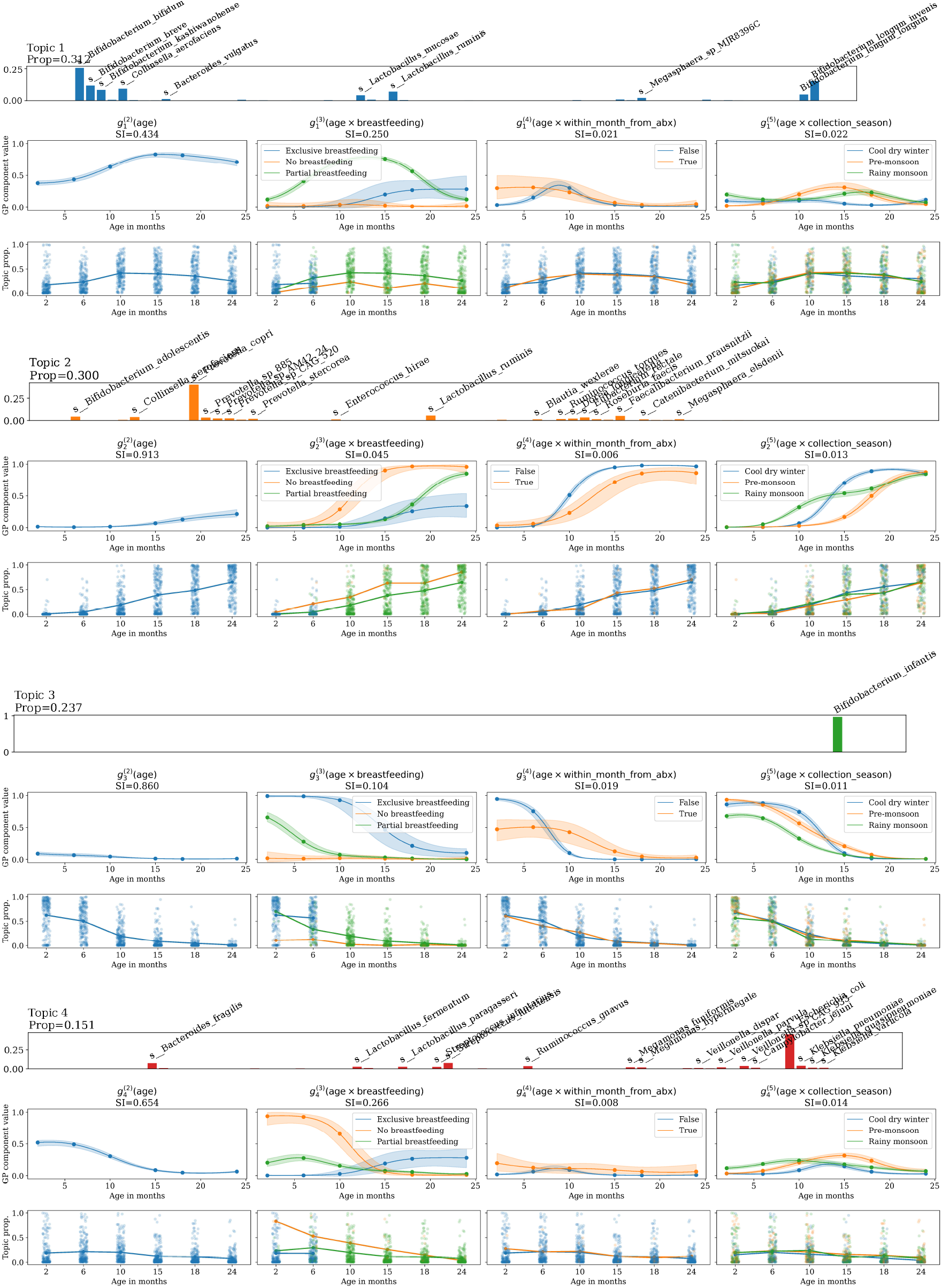
Visualization of topics, latent functions, and topic proportions of samples in Dhaka dataset (*L* = 4).

## References

[1] T.S. Schmidt, J. Raes, and P. Bork, The human gut microbiome: from association to modulation, Cell 172 (2018), pp. 1198–1215.

[2] C.J. Stewart, N.J. Ajami, J.L. O’Brien, D.S. Hutchinson, D.P. Smith, M.C. Wong, M.C. Ross, R.E. Lloyd, H. Doddapaneni, G.A. Metcalf, et al., Temporal development of the gut microbiome in early childhood from the teddy study, Nature 562 (2018), pp. 583–588.

[3] H. Peng, S. Andreu-Sanchez, A.J. Ruiz-Moreno, A. Fernández-Pato, J. Wu, R. Gacesa, A. Zhernakova, D. Wang, and J. Fu, Longitudinal gut microbiota tracking reveals the dynamics of horizontal gene transfer, Nature Communications (2025).

[4] L.A. David, A.C. Materna, J. Friedman, M.I. Campos-Baptista, M.C. Blackburn, A. Perrotta, S.E. Erdman, and E.J. Alm, Host lifestyle affects human microbiota on daily timescales, Genome biology 15 (2014), p. R89.

[5] L. Shenhav, O. Furman, L. Briscoe, M. Thompson, J.D. Silverman, I. Mizrahi, and E. Halperin, Modeling the temporal dynamics of the gut microbial community in adults and infants, PLoS computational biology 15 (2019), p. e1006960.

[6] L. Grieneisen, R. Blekhman, and E. Archie, How longitudinal data can contribute to our understanding of host genetic effects on the gut microbiome, Gut Microbes 15 (2023), p. 2178797.

[7] X. Zhou, X. Shen, J.S. Johnson, D.J. Spakowicz, M. Agnello, W. Zhou, M. Avina, A. Honkala, F. Chleilat, S.J. Chen, et al., Longitudinal profiling of the microbiome at four body sites reveals core stability and individualized dynamics during health and disease, Cell Host & Microbe 32 (2024), pp. 506–526.

[8] H. Mallick, A. Rahnavard, L.J. McIver, S. Ma, Y. Zhang, L.H. Nguyen, T.L. Tickle, G. Weingart, B. Ren, E.H. Schwager, et al., Multivariable association discovery in population-scale meta-omics studies, PLoS computational biology 17 (2021), p. e1009442.

[9] V. Bucci, B. Tzen, N. Li, M. Simmons, T. Tanoue, E. Bogart, L. Deng, V. Yeliseyev, M.L. Delaney, Q. Liu, et al., Mdsine: Microbial dynamical systems inference engine for microbiome time-series analyses, Genome biology 17 (2016), p. 121.

[10] T.A. Joseph, L. Shenhav, J.B. Xavier, E. Halperin, and I. Pe’er, Compositional lotka-volterra describes microbial dynamics in the simplex, PLoS computational biology 16 (2020), p. e1007917.

[11] T. Äijö, C.L. Müller, and R. Bonneau, Temporal probabilistic modeling of bacterial compositions derived from 16s rrna sequencing, Bioinformatics 34 (2018), pp. 372–380.

[12] L. Cheng, S. Ramchandran, T. Vatanen, N. Lietzén, R. Lahesmaa, A. Vehtari, and H. Lähdesmäki, An additive gaussian process regression model for interpretable non-parametric analysis of longitudinal data, Nature communications 10 (2019), p. 1798.

[13] W. Cao, D. Wang, J. Li, H. Zhou, L. Li, and Y. Li, Brits: Bidirectional recurrent imputation for time series, Advances in neural information processing systems 31 (2018).

[14] Y. Luo, X. Cai, Y. Zhang, J. Xu, et al., Multivariate time series imputation with generative adversarial networks, Advances in neural information processing systems 31 (2018).

[15] W. Du, D. Côté, and Y. Liu, Saits: Self-attention-based imputation for time series, Expert Systems with Applications 219 (2023), p. 119619.

[16] J.M. Choi, M. Ji, L.T. Watson, and L. Zhang, Deepmicrogen: a generative adversarial network-based method for longitudinal microbiome data imputation, Bioinformatics 39 (2023), p. btad286.

[17] E. De Brouwer, J. Simm, A. Arany, and Y. Moreau, Gru-ode-bayes: Continuous modeling of sporadically-observed time series, Advances in neural information processing systems 32 (2019).

[18] Y. Qu, R. Lyu, D. Wang, Y. Dai, A. Turcan, S. Yu, J. Xie, J. Roach, C. Butler, P.T. Yap, et al., Deep-learning-based interpolation of longitudinal microbiome data powers biologically informative discovery, bioRxiv (2025), pp. 2025–02.

[19] D.P. Kingma and M. Welling, Auto-encoding variational bayes, arXiv preprint arXiv:1312.6114 (2013).

[20] K. Sohn, H. Lee, and X. Yan, Learning structured output representation using deep conditional generative models, Advances in neural information processing systems 28 (2015).

[21] F.P. Casale, A. Dalca, L. Saglietti, J. Listgarten, and N. Fusi, Gaussian process prior variational autoencoders, Advances in neural information processing systems 31 (2018).

[22] V. Fortuin, D. Baranchuk, G. Rätsch, and S. Mandt, Gp-vae: Deep probabilistic time series imputation, in International conference on artificial intelligence and statistics. PMLR, 2020, pp. 1651–1661.

[23] S. Ramchandran, G. Tikhonov, K. Kujanpää, M. Koskinen, and H. Lähdesmäki, Longitudinal variational autoencoder, in International Conference on Artificial Intelligence and Statistics. PMLR, 2021, pp. 3898–3906.

[24] M.Y. Balık, M. Sinelnikov, P. Ong, and H. Lähdesmäki, Bayesian Basis Function Approximation for Scalable Gaussian Process Priors in Deep Generative Models, in Forty-second International Conference on Machine Learning. 2025.

[25] D.M. Blei, A.Y. Ng, and M.I. Jordan, Latent dirichlet allocation, Journal of machine Learning research 3 (2003), pp. 993–1022.

[26] D. Lee and H.S. Seung, Algorithms for non-negative matrix factorization, Advances in neural information processing systems 13 (2000).

[27] A. Srivastava and C. Sutton, Autoencoding Variational Inference For Topic Models, in International Conference on Learning Representations. 2017.

[28] Y. Miao, E. Grefenstette, and P. Blunsom, Discovering discrete latent topics with neural variational inference, in International conference on machine learning. PMLR, 2017, pp. 2410–2419.

[29] A.B. Dieng, F.J. Ruiz, and D.M. Blei, Topic modeling in embedding spaces, Transactions of the Association for Computational Linguistics 8 (2020), pp. 439–453.

[30] Y. Zhao, H. Cai, Z. Zhang, J. Tang, and Y. Li, Learning interpretable cellular and gene signature embeddings from single-cell transcriptomic data, Nature communications 12 (2021), p. 5261.

[31] A.W. Lynch, C.V. Theodoris, H.W. Long, M. Brown, X.S. Liu, and C.A. Meyer, Mira: joint regulatory modeling of multimodal expression and chromatin accessibility in single cells, Nature methods 19 (2022), pp. 1097–1108.

[32] S. Hosoda, S. Nishijima, T. Fukunaga, M. Hattori, and M. Hamada, Revealing the microbial assemblage structure in the human gut microbiome using latent dirichlet allocation, Microbiome 8 (2020), p. 95.

[33] C. Frioux, R. Ansorge, E. Özkurt, C.G. Nedjad, J. Fritscher, C. Quince, S.M. Waszak, and F. Hildebrand, Enterosignatures define common bacterial guilds in the human gut microbiome, Cell Host & Microbe 31 (2023), pp. 1111–1125.

[34] P. Jeganathan and S.P. Holmes, A statistical perspective on the challenges in molecular microbial biology, Journal of Agricultural, Biological and Environmental Statistics 26 (2021), pp. 131–160.

[35] K. Sankaran and S.P. Holmes, Latent variable modeling for the microbiome, Biostatistics 20 (2019), pp. 599–614.

[36] J. Fukuyama, L. Rumker, K. Sankaran, P. Jeganathan, L. Dethlefsen, D.A. Relman, and S.P. Holmes, Multidomain analyses of a longitudinal human microbiome intestinal cleanout perturbation experiment, PLoS computational biology 13 (2017), p. e1005706.

[37] J. Fukuyama, K. Sankaran, and L. Symul, Multiscale analysis of count data through topic alignment, Biostatistics 24 (2023), pp. 1045–1065.

[38] C.K. Williams and C.E. Rasmussen, Gaussian processes for machine learning, Vol. 2, MIT press Cambridge, MA, 2006.

[39] A. Solin and S. Särkkä, Hilbert space methods for reduced-rank gaussian process regression, Statistics and Computing 30 (2020), pp. 419–446.

[40] J. Timonen and H. Lähdesmäki, Scalable mixed-domain gaussian process modeling and model reduction for longitudinal data, Bayesian Analysis 1 (2025), pp. 1–28.

[41] D.T. Truong, E.A. Franzosa, T.L. Tickle, M. Scholz, G. Weingart, E. Pasolli, A. Tett, C. Huttenhower, and N. Segata, Metaphlan2 for enhanced metagenomic taxonomic profiling, Nature methods 12 (2015), pp. 902–903.

[42] S. Nishijima, E. Stankevic, O. Aasmets, T.S. Schmidt, N. Nagata, M.I. Keller, P. Ferretti, H.B. Juel, A. Fullam, S.M. Robbani, et al., Fecal microbial load is a major determinant of gut microbiome variation and a confounder for disease associations, Cell 188 (2025), pp. 222–236.

[43] M.A. Alvarez, L. Rosasco, N.D. Lawrence, et al., Kernels for vector-valued functions: A review, Foundations and Trends® in Machine Learning 4 (2012), pp. 195–266.

[44] C. Boutsidis and E. Gallopoulos, Svd based initialization: A head start for nonnegative matrix factorization, Pattern recognition 41 (2008), pp. 1350–1362.

[45] D.P. Kingma, Adam: A method for stochastic optimization, arXiv preprint arXiv:1412.6980 (2014).

[46] J. Yan, G. Chuai, T. Qi, F. Shao, C. Zhou, C. Zhu, J. Yang, Y. Yu, C. Shi, N. Kang, et al., Metatopics: an integration tool to analyze microbial community profile by topic model, BMC genomics 18 (2017), p. 962.

[47] R. Deveaud, E. SanJuan, and P. Bellot, Accurate and effective latent concept modeling for ad hoc information retrieval, Document numérique 17 (2014), pp. 61–84.

[48] M. Zhu and A. Ghodsi, Automatic dimensionality selection from the scree plot via the use of profile likelihood, Computational Statistics & Data Analysis 51 (2006), pp. 918–930.

[49] I.M. Sobol, Global sensitivity indices for nonlinear mathematical models and their monte carlo estimates, Mathematics and computers in simulation 55 (2001), pp. 271–280.

[50] A. Saltelli, P. Annoni, I. Azzini, F. Campolongo, M. Ratto, and S. Tarantola, Variance based sensitivity analysis of model output. design and estimator for the total sensitivity index, Computer physics communications 181 (2010), pp. 259–270.

[51] F. Gamboa, A. Janon, T. Klein, and A. Lagnoux, Sensitivity indices for multi-variate outputs, Comptes Rendus. Mathématique 351 (2013), pp. 307–310.

[52] T. Vatanen, Q.Y. Ang, L. Siegwald, S.A. Sarker, C.I. Le Roy, S. Duboux, O. Delannoy-Bruno, C. Ngom-Bru, C.L. Boulangé, M. Stražar, et al., A distinct clade of bifidobacterium longum in the gut of bangladeshi children thrives during weaning, Cell 185 (2022), pp. 4280–4297.

[53] T. Vatanen, D.R. Plichta, J. Somani, P.C. Münch, T.D. Arthur, A.B. Hall, S. Rudolf, E.J. Oakeley, X. Ke, R.A. Young, et al., Genomic variation and strain-specific functional adaptation in the human gut microbiome during early life, Nature microbiology 4 (2019), pp. 470–479.

[54] J. Lloyd-Price, C. Arze, A.N. Ananthakrishnan, M. Schirmer, J. Avila-Pacheco, T.W. Poon, E. Andrews, N.J. Ajami, K.S. Bonham, C.J. Brislawn, et al., Multiomics of the gut microbial ecosystem in inflammatory bowel diseases, Nature 569 (2019), pp. 655–662.

[55] V. Pawlowsky-Glahn, J.J. Egozcue, and R. Tolosana-Delgado, Modeling and analysis of compositional data, John Wiley & Sons, 2015.

[56] J.A. Martín-Fernández, C. Barceló-Vidal, and V. Pawlowsky-Glahn, Dealing with zeros and missing values in compositional data sets using nonparametric imputation, Mathematical Geology 35 (2003), pp. 253–278.

[57] D.M. Endres and J.E. Schindelin, A new metric for probability distributions, IEEE Transactions on Information theory 49 (2003), pp. 1858–1860.

[58] D.F. Crouse, On implementing 2d rectangular assignment algorithms, IEEE Transactions on Aerospace and Electronic Systems 52 (2016), pp. 1679–1696.

[59] M. Modesto, C. Ngom-Bru, D. Scarafile, A. Bruttin, S. Pruvost, S.A. Sarker, T. Ahmed, O. Sakwinska, P. Mattarelli, and S. Duboux, Bifidobacterium longum subsp. iuvenis subsp. nov., a novel subspecies isolated from the faeces of weaning infants, International Journal of Systematic and Evolutionary Microbiology 73 (2023), p. 006013.

[60] T. Vatanen, A.D. Kostic, E. d’Hennezel, H. Siljander, E.A. Franzosa, M. Yassour, R. Kolde, H. Vlamakis, T.D. Arthur, A.M. Hämäläinen, et al., Variation in microbiome lps immunogenicity contributes to autoimmunity in humans, Cell 165 (2016), pp. 842–853.

[61] Y. Shao, S.C. Forster, E. Tsaliki, K. Vervier, A. Strang, N. Simpson, N. Kumar, M.D. Stares, A. Rodger, P. Brocklehurst, et al., Stunted microbiota and opportunistic pathogen colonization in caesarean-section birth, Nature 574 (2019), pp. 117–121.

[62] H. Nunez, P.A. Nieto, R.A. Mars, M. Ghavami, C. Sew Hoy, and K. Sukhum, Early life gut microbiome and its impact on childhood health and chronic conditions, Gut microbes 17 (2025), p. 2463567.

[63] Y. Yan, Y. Lei, Y. Qu, Z. Fan, T. Zhang, Y. Xu, Q. Du, D. Brugger, Y. Chen, K. Zhang, et al., Bacteroides uniformis-induced perturbations in colonic microbiota and bile acid levels inhibit th17 differentiation and ameliorate colitis developments, npj Biofilms and Microbiomes 9 (2023), p. 56.

[64] D. Ishikawa, X. Zhang, K. Nomura, T. Shibuya, M. Hojo, M. Yamashita, S. Koizumi, F. Yamazaki, S. Iwamoto, M. Saito, et al., Anti-inflammatory effects of bacteroidota strains derived from outstanding donors of fecal microbiota transplantation for the treatment of ulcerative colitis, Inflammatory Bowel Diseases 30 (2024), pp. 2136–2145.

[65] J. Timonen, H. Mannerström, A. Vehtari, and H. Lähdesmäki, lgpr: an interpretable non-parametric method for inferring covariate effects from longitudinal data, Bioinformatics 37 (2021), pp. 1860–1867.

[66] T. Wang and H. Zhao, A dirichlet-tree multinomial regression model for associating dietary nutrients with gut microorganisms, Biometrics 73 (2017), pp. 792–801.

[67] F. Beghini, L.J. McIver, A. Blanco-Míguez, L. Dubois, F. Asnicar, S. Maharjan, A. Mailyan, P. Manghi, M. Scholz, A.M. Thomas, et al., Integrating taxonomic, functional, and strain-level profiling of diverse microbial communities with biobakery 3, elife 10 (2021), p. e65088.

[68] F. Meyer, D. Paarmann, M. D’Souza, R. Olson, E.M. Glass, M. Kubal, T. Paczian, A. Rodriguez, R. Stevens, A. Wilke, et al., The metagenomics rast server–a public resource for the automatic phylogenetic and functional analysis of metagenomes, BMC bioinformatics 9 (2008), p. 386.

[69] G.M. Douglas, V.J. Maffei, J.R. Zaneveld, S.N. Yurgel, J.R. Brown, C.M. Taylor, C. Huttenhower, and M.G. Langille, Picrust2 for prediction of metagenome functions, Nature biotechnology 38 (2020), pp. 685–688.

[70] A. Paszke, S. Gross, F. Massa, A. Lerer, J. Bradbury, G. Chanan, T. Killeen, Z. Lin, N. Gimelshein, L. Antiga, et al., Pytorch: An imperative style, high-performance deep learning library, Advances in neural information processing systems 32 (2019).

[71] W. Du, Pypots: a python toolbox for data mining on partially-observed time series, arXiv preprint arXiv:2305.18811 (2023).

[72] T. Akiba, S. Sano, T. Yanase, T. Ohta, and M. Koyama, Optuna: A next-generation hyperparameter optimization framework, in Proceedings of the 25th ACM SIGKDD international conference on knowledge discovery & data mining. 2019, pp. 2623–2631.

[73] F. Pedregosa, G. Varoquaux, A. Gramfort, V. Michel, B. Thirion, O. Grisel, M. Blondel, P. Prettenhofer, R. Weiss, V. Dubourg, et al., Scikit-learn: Machine learning in python, the Journal of machine Learning research 12 (2011), pp. 2825–2830.

